# SAP loss limits anti-insulin atypical B cell activation and pro-inflammatory CD8 T cells despite preserved Tfh responses to protect against type 1 diabetes

**DOI:** 10.64898/2026.07.29.741363

**Authors:** Landon M. Clark, Dudley H. McNitt, Jack C. McAninch, Lindsay E. Bass, Marguerite L. Padgett, Andres F. Moreno, Christina T. Brannon, Casey M. Nichols, Matthew T. Stier, Rachel H. Bonami

## Abstract

SLAM-associated protein (SAP) is required for T follicular helper (Tfh)-B cell interactions that underlie germinal center formation, but it is unclear if SAP governs islet-reactive CD4+ T cell-B cell interactions and downstream pro-inflammatory CD8+ T cell destruction of islets in type 1 diabetes (T1D). To address this question, we utilized the VH125^SD^.NOD mouse model, whereby 1-3% of all B cells bind insulin. Germline *SAP* loss in this model led to reduced T1D incidence and impaired germinal center B cell formation, yet did not alter T follicular helper cell formation or phenotype. *SAP* loss reduced pro-inflammatory and activated insulin-autoreactive B-T interactions and limited anti-insulin B cell proliferation, activation, and upregulation of co-stimulatory molecules otherwise enhanced in the pancreas. Anti-insulin extrafollicular antibody and memory responses following immunization were preserved in VH125^SD^.*SAP^-/-^*.NOD mice, but activated atypical anti-insulin B cell responses were reduced. Ultimately, *SAP* loss led to reduced pro-inflammatory CD8+ T cell formation and islet-reactive progenitor exhausted CD8+ T cells in pancreata. These data highlight the essential role of SAP in mediating proinflammatory, anti-insulin B-T interactions to support T1D.

**Graphical Abstract:** 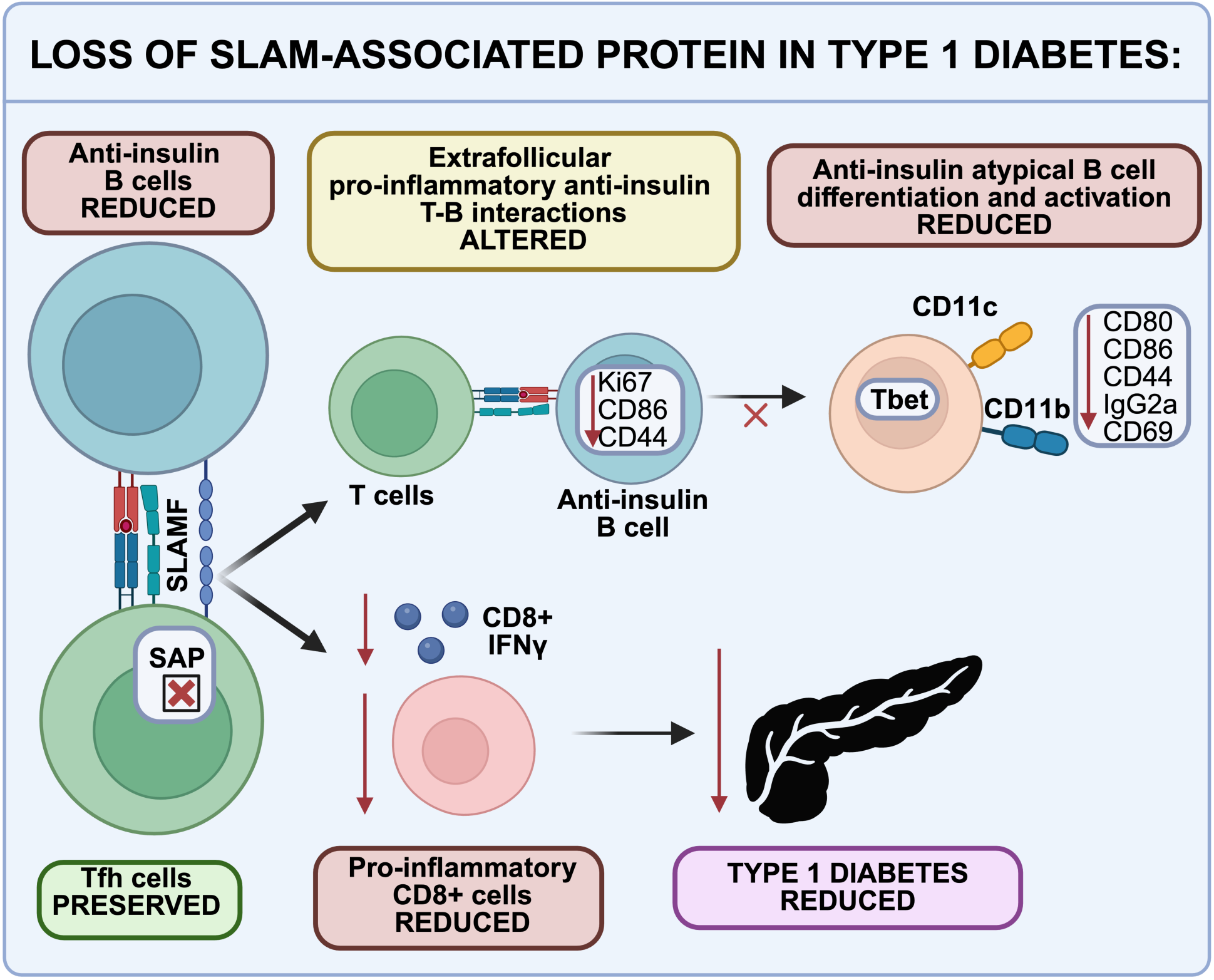

## Introduction

Autoreactive CD4+ T-B interactions are thought to license pathogenic CD8+ T cells to destroy beta-cells in T1D^1^. In line with this, teplizumab (anti-CD3) is an approved immunotherapy for stage 2 and stage 3 T1D, which broadly impacts T cell function to delay, but not prevent diabetes onset^2–4^. The B lymphocyte depleting agent, rituximab (anti-CD20), was also linked with temporarily preserving beta cell function in a clinical trial but was associated with concomitant vaccine response impairment^5,6^. Autoreactive B cells in T1D target the key autoantigen, insulin, with insulin autoantibodies (signaling pathologic anti-insulin B/T lymphocyte interactions) predicting diabetes in mice and humans^7^. NOD mice which lack insulin-binding B cells (but retain other specificities) are protected against diabetes^8^, and B cells are chiefly thought to drive T1D by presenting antigen to T cells^9–11^. Thus, understanding the mechanisms underlying pro-pathogenic T-B lymphocyte interactions holds promise for advancing T1D immunotherapies.

T-B lymphocyte interactions occur via germinal center (GC) or extrafollicular (EF) responses^12,13^. GCs facilitate long-lived immunity whereby B cells undergo somatic hypermutation and affinity maturation via Tfh interactions to generate plasma cells and memory B cells^14^. Alternatively, B cells can enter an EF response to generate atypical (age-associated) B cells, memory B cells, and plasmablasts, with atypical B cell responses noted in settings of chronic infection, autoimmunity, and aging^12,15,16^. The term “atypical B cells” is a broad category encompassing several distinct populations, with core markers including CD11c, CD11b, and/or T-bet in mice, whereas “EF responses” refers to the specific anatomical niche in which they form^12^. Presently, the role for atypical B cell:T cell interactions in the EF niche to support T1D pathogenesis is unclear. T cells depend on the transcriptional repressor BCL6 to support both GC and EF responses^17,18^. By comparison, SLAM-associated protein (SAP) is required for GC T-B responses but dispensable for EF responses, although its role regarding atypical B cell formation is still unclear^19–21^. SAP deficiency could therefore enable the study of how T-B cell interactions contribute to T1D in the absence of GCs.

SAP is essential for Tfh interactions with B cells^20^. In T1D, Tfh/Tph populations are elevated in individuals, and adoptively transferred Tfh cells elicit diabetes in NOD.RAG recipients^22,23^. Our recent data show that T cell loss of the key GC protein, BCL6, blunts Tfh and GC B cell differentiation and blocks diabetes development in NOD mice^24^. The specific mechanisms underlying this diabetes protection are unclear, as T cell loss of BCL6 impacted both GC and atypical B cell EF responses^18,25^. It is thus possible that Tfh-like T cell interactions with autoreactive B cells outside of GCs are contributing to T1D pathogenesis. To investigate anti-insulin B cell activation and proliferation in diabetes in a setting with abolished GCs but preserved EF responses, we generated VH125^SD^.SAP^-/-^.NOD (VH125^SD^ΔSAP) mice, as they should have a trackable anti-insulin B cell population (unlike wildtype (WT) NOD mice)^26^. Specifically, the VH125^SD^.NOD model has an IgH locus-targeted B cell receptor heavy chain transgene which generates a 1-3% population of anti-insulin B cells amongst the total B cell repertoire^27^.

Here, we show that VH125^SD^ΔSAP NOD mice have few GC B cells but persistent Tfh-like cells that are phenotypically similar to their VH125^SD^.NOD counterparts. Despite this preserved Tfh pool, diabetes was significantly reduced in the SAP-deficient mice. This protection was associated with impaired autoreactive CD4+ T-B interactions, as demonstrated by reduced non-GC CD44 and CCR6 expression in CD4+ T-anti-insulin B cell conjugates. SAP loss reduced non-GC anti-insulin B cell upregulation of activation, co-stimulatory, and proliferative markers. Immunization studies revealed that loss of SAP blocked anti-insulin, atypical B cell expansion (T-bet+ CD11c+ and CD11b+ CD11c+), with both populations expressing increased CD44, CD69, CD86, CD80, and were IgG2a+ switched compared to other anti-insulin B cell subsets. Loss of SAP reduced several pro-inflammatory CD8+ T cell populations, including islet-reactive T cell progenitor exhausted (Tpex), CXCR3+, and IFN-γ+ cells in the pancreatic lymph nodes (pLNs) and pancreas of SAP-deficient VH125^SD^.NOD mice. This work provides evidence that disrupting key T-B cognate interactions could alter diabetes development even when immune tolerance for insulin fails to prevent peripheral accumulation of anti-insulin B lymphocytes.

## Methods

### Sex as a biological variable

All studies used female mice, in line with increased diabetes incidence in females relative to males in NOD mice^28^.

### Animals

*SAP-*deficient mice on the C57BL/6 background were provided by Dr. Pamela Schwartzberg (National Institutes of Health), bred in our colony, and subsequently backcrossed into the NOD strain as described previously^29^. Anti-insulin VH125^SD^.NOD BCR transgenic mice were described previously^27^. SAP-deficient NOD mice were then crossed with VH125^SD^.NOD mice to generate SAP-deficient VH125^SD^.NOD mice. All mice were housed under specific pathogen–free conditions, littermate controls were used for diabetes monitoring, while mice used for other experiments were co-housed for a minimum of 2 weeks prior to use. All animal studies were approved by the fully AAALAC-accredited Vanderbilt University Institutional Animal Care and Use Committee.

### Diabetes monitoring

Blood glucose was measured weekly via a glucometer from 10-40 weeks of age by nicking tails. Diabetes was diagnosed after two consecutive blood glucose readings >250 mg/dL.

### Histological assessment of insulitis and tertiary lymphoid structure organization

Mice were sacrificed and pancreata were dissected from non-diabetic female mice. Pancreata were incubated overnight in 10% formalin at room temperature and then dehydrated briefly in 70% ethanol and paraffin-embedded by the Vanderbilt Tissue Pathology Shared Resource (TPSR). 10 µm sections were cut, deparaffinized, and subsequently stained with H&E by TPSR staff. Slide images were obtained using a bright field Aperio ScanScope CS. Visualization and analysis was performed by using ImageScope software (Leica Biosystems). Islets were individually blind scored for insulitis severity (lymphocyte invasion), ranging from 0 (no insulitis), 1 (1-25%), 2 (26-50%), 3 (51-75%), and 4 (>75%) of the islet area occupied by immune infiltrate.

Pancreas blocks exhibiting grade 3-4 insulitis (as scored above) were chosen from n = 6 mice per genotype for subsequent CD3 and B220 IHC staining using a Leica Bond-Max IHC stainer. Following heat-induced antigen retrieval with Bond Epitope Retrieval Solution 2 (Leica Biosystems), 10 µm pancreas sections were stained by TPSR core staff with antibodies reactive against either CD45R/B220 (RA3–6B2) or CD3 (M-20). Islets were blind scored for average infiltration of CD3 (T cells) and B220 (B cells), using a scoring scale of 0 (no infiltrate), 1 (peri-insulitis), 2 (mild lymphocytic infiltrate) or 3 (moderate/severe lymphocytic infiltrate). Islets with an insulitis severity score of 3 or higher were further blind scored as “organized” (defined by organized B/T cell zones) or having “non-discrete zones” (defined by B/T infiltration that was not organized into discrete B or T cell zones).

### Cell isolation from tissues

Cells were isolated from spleen, pLNs, and pancreas, as previously described^26^. In brief, spleens were freshly harvested and macerated through a 70-µm cell strainer with either Hank’s balanced salt solution (HBSS) with 10% bovine calf serum or FACS buffer (1x PBS + 1 mmol/L EDTA + 5% FBS). Cells were pelleted, red blood cells were lysed with red blood cell lysis solution (140 mmol/L NH_4_Cl + 17 mmol/L Tris), and cells were resuspended in FACS buffer; lymph nodes were processed similarly but without red blood cell lysis. Pancreata were digested with 1 mg/mL of collagenase *P* (Millipore-Sigma #11249002001) in HBSS and incubated while shaking at 37°C for 10 minutes, then tissue was disrupted with an 18G needle. HBSS+10% bovine calf serum was immediately added to inhibit collagenase activity. Cells were filtered through a 70-µm cell strainer, washed in HBSS + 10% bovine calf serum, pelleted, and resuspended in FACS buffer. Cell counts were performed for each organ prior to downstream flow cytometry staining and analysis, using a hemocytometer (pancreata) or Countess Automated Cell Counter (ThermoFisher, lymph nodes, spleen).

Cells were stained for flow cytometry, with murine reactive antibodies listed in **Supplemental Table 1.** Biotinylated human insulin (Millipore-Sigma #I2643) was generated and used to detect insulin-binding specificity, whereas insulin-occupied BCRs were detected via biotinylated anti-insulin mAb123 (HB-123; American Type Culture Collection) as described previously^8,25,30^.

Cells were incubated with Fc Block (BD Biosciences) (and non-biotinylated human insulin for pancreata), followed by surface Ab, tetramer (NRP-V7, insulin B chain fragment, NIH Tetramer Core Facility, **Supplemental Table 1**), and/or biotinylated human insulin or mAb123 for 1 hour at 4°C in FACS buffer (detected by streptavidin-fluorochrome). Cells were then incubated with BD transcription factor staining set (BD Pharmingen, Cat# 562574) for 30 minutes at 4°C/on ice, then stained with the intracellular Ab mix for 1 hour or overnight at 4°C/on ice. After overnight incubation, cells were washed three times with 1x permeabilization buffer, resuspended in FACS buffer, and acquired on a BD Fortessa or Cytek Aurora cytometers.

### Insulin-Brucella abortus ring test antigen (BRT) immunization

Recombinant human insulin (Sigma-Aldrich #I2643) was conjugated to *Brucella abortus* ring test antigen (BRT; U.S. Department of Agriculture Animal and Plant Health Inspection Services, Ames, IA) as previously described^31^. Briefly, recombinant human insulin was dissolved in 0.1M Bicine, mixed with M-maleimidobenzoyl-N-hydroxy-succinimide in dimethylformamide) and incubated for 2 hours at room temperature with gentle agitation. Functionalized insulin was precipitated out using cold 0.1 M citrate phosphate buffer and dissolved in 0.02 M bicine saline. BRT solution was washed three times with bicine saline containing dithiothreitol until supernatant was clear. BRT was resuspended in 1 mL of bicine saline with dithiothreitol incubated with 2-iminothiolane at room temperature for 10 minutes with gentle agitation. BRT was washed with bicine saline, resuspended in the functionalized insulin bicine saline solution, and incubated at room temperature for 2 hours then overnight at 4°C. The insulin-BRT solution was spun down, and supernatant was removed to remove free insulin and then resuspended in 1x PBS. VH125^SD^ and VH125^SD^.SAP^-/-^.NOD mice were immunized with insulin-BRT or unconjugated BRT i.p. and spleen harvested 5-7 days after immunization. Sera was collected both pre-immunization (Day 0) and post-immunization (Days 5-7).

### ELISA

Competitive binding ELISA was used to measure insulin-specific antibodies. Specifically, 384-well Maxisorp Nunc plates (Thermo Scientific) were coated with 1 μg/ml human insulin (Sigma-Aldrich #I2643) in borate-buffered saline overnight at 37°C. Plates were washed 5X with 0.5X PBS, blocked with 2% bovine sera albumin in 1X PBS, then washed 5X with 0.5X PBS. Sera (isolated as in ^25^) were diluted 1:100 in 1X PBS+0.1% Tween-20 (PBS-T) and incubated in wells for 1 hour at room temperature. To measure insulin-specific IgG, parallel samples were incubated in the presence of 100 μg/ml human insulin in PBS-T to enable subtraction of these O.D. values from non-inhibited well O.D.s. After sera incubation, plates were washed 5X with 0.5X PBS, and incubated with goat anti mouse IgM-alkaline phosphatase (AP) (Southern Biotech 1020-04), IgG1-AP (Southern Biotech 1070-04), IgG2a-AP (Southern Biotech 1080-04), IgG2b-AP (Southern Biotech 1090-04), and IgG3-AP (Southern Biotech 1100-04) diluted 1:2000 in TBS+0.1% Tween-20 in independent wells to detect isotype specific anti-insulin antibodies. After 1 hour incubation, plates were washed 10X with 0.5X PBS. Wells were incubated with 10 mg/ml p-nitrophenyl phosphate substrate (Sigma-Aldrich) in a 50 mM potassium carbonate + 1 mM magnesium chloride buffer. Optical density was read at 405 nm after 1 hour using a Synergy LX Microplate Autoreader (Bio-Tek). Isotype controls coated at 1 µg/mL were used to confirm specificity of AP conjugated detection antibodies (mouse IgM, Invitrogen #02-6800, mouse IgG1, Invitrogen #02-6100; mouse IgG2a, Invitrogen #02-6200; mouse IgG2b Invitrogen #02-6300, mouse IgG3, #MA1-10433).

### Cytokine stimulation

For intracellular cytokine staining, cells were isolated from spleen and pancreatic draining lymph nodes as above. For cells within pancreas, pancreas was processed as above but resuspended in MACS buffer and T cells positively selected using a Miltenyi CD3ε MicroBead Kit (Cat# 130-094-973). CD3+ cells were resuspended in Iscove’s Modified Dulbecco’s Medium (10% Fetal Bovine Serum, 2 mM L-glutamine, 5 mM sodium pyruvate, 1x MEM non-essential amino acids, 10 mM HEPES, 100 U/mL penicillin/streptomycin, 20 µM β-mercaptoethanol) at 5-10×10^6^ cells/mL. Cells were incubated for 2 hours at 37°C with or without stimulation (500 ng/mL PMA + 5 ug/mL Brefeldin A + 1.3 µM ionomycin), washed 3 times, then resuspended in FACS buffer and stained as above with modifications; after surface staining for 1 hour, cells were fixed for 20 minutes with 2% paraformaldehyde in 1x PBS on ice, then incubated with eBioscience intracellular fixation & permeabilization buffer (Cat#: 88-8824-00) for 5 minutes at room temperature. Intracellular staining for intracellular cytokines was done for 1 hour on ice for primary antibodies, and 30 minutes on ice for secondary antibodies (**Supplemental Table 1**).

### Flow cytometry analysis and unsupervised clustering

Flow cytometry was analyzed via FlowJo v10 and Prism v10 for representative gating and graphs. For minimally supervised analysis, files were imported into OMIQ (Dotmatics) and scaled using the arcsinh cofactor 10, followed by gating on the insulin+ B220+ CD19+ CD45+ population and subsampling to ∼5000 cells per sample, ∼100,000 cells total. Normalization was performed via CyCombine software^32^, followed by opt-SNE analysis^33^. Phenograph (k=20, Annoy algorithm, Euclidean distance metric, Leiden clustering, seed 2730) was used to create initial clustering^34^, followed by heatmap analysis and manual canonical gating to combine or separate further clusters as necessary. Diffcyt-edgeR-DA (differential abundance)^35^ was used both in the initial Phenograph clustering analysis and the combined clustering analysis to identify regions of |log2FC| > 1 with p value < 0.01^35^.

### Statistical analyses

Statistical tests are indicated in the corresponding figure legends and significance values were calculated using GraphPad Prism v9.3.1 (GraphPad Software).

### Study approval

All experiments involving animals were approved by Vanderbilt University IACUC, Animal protocol Ethics Approval Number #M1800152

## Data availability

Values for individual data points are indicated in graphs. Raw data files are available upon reasonable request.

## Results

### Loss of SAP impairs anti-insulin B cell infiltration of the pancreas and germinal center B cell formation in insulin-skewed VH125^SD^ NOD mice

To clarify the role of GC vs. EF responses in supporting anti-insulin T-B communication and downstream pathology, we generated VH125^SD^ NOD and VH125^SD^ΔSAP NOD mice as in Methods, in which an anti-insulin, IgH-targeted BCR transgene supports the formation of a 1-3% population of anti-insulin B cells^27^ (**Figure 1A)**. Accelerated diabetes occurs in the VH125^SD^ NOD model compared to the NOD model^27^. In another autoimmune disease, SAP loss abolished GC responses while EF plasmablast responses remained intact^21^. Thus, the VH125^SD^ΔSAP NOD model provides a means to study how disrupting anti-insulin B/T lymphocyte GC responses impacts diabetes pathogenesis while preserving their ability to interact in the EF niche. We first examined whether loss of SAP impacted (1) anti-insulin B cell infiltration of the pancreas and (2) spontaneous GC B cell formation in a T1D context. Anti-insulin B cells proportions were similar in the spleen but were reduced by SAP loss in the pLNs and pancreas (**Figure 1B**). Proportions and total numbers of CD4+ T cells, CD8+ T cells, and CD19+ B cells were unchanged in all organs, except for a small increase in the proportion (but not number) of CD8+ T cells in the pLNs **(Supplemental Figure 1A-F)**. GC B cell frequencies were significantly reduced by SAP loss in both pLNs and pancreas in VH125^SD^.NOD mice, with GC B cell numbers also reduced in pLNs, as expected (**Figure 1C**). Together, these results highlight that anti-insulin B cell infiltration and GC formation is reduced by loss of SAP in insulin-skewed VH125^SD^.NOD mice.

**Figure 1:**
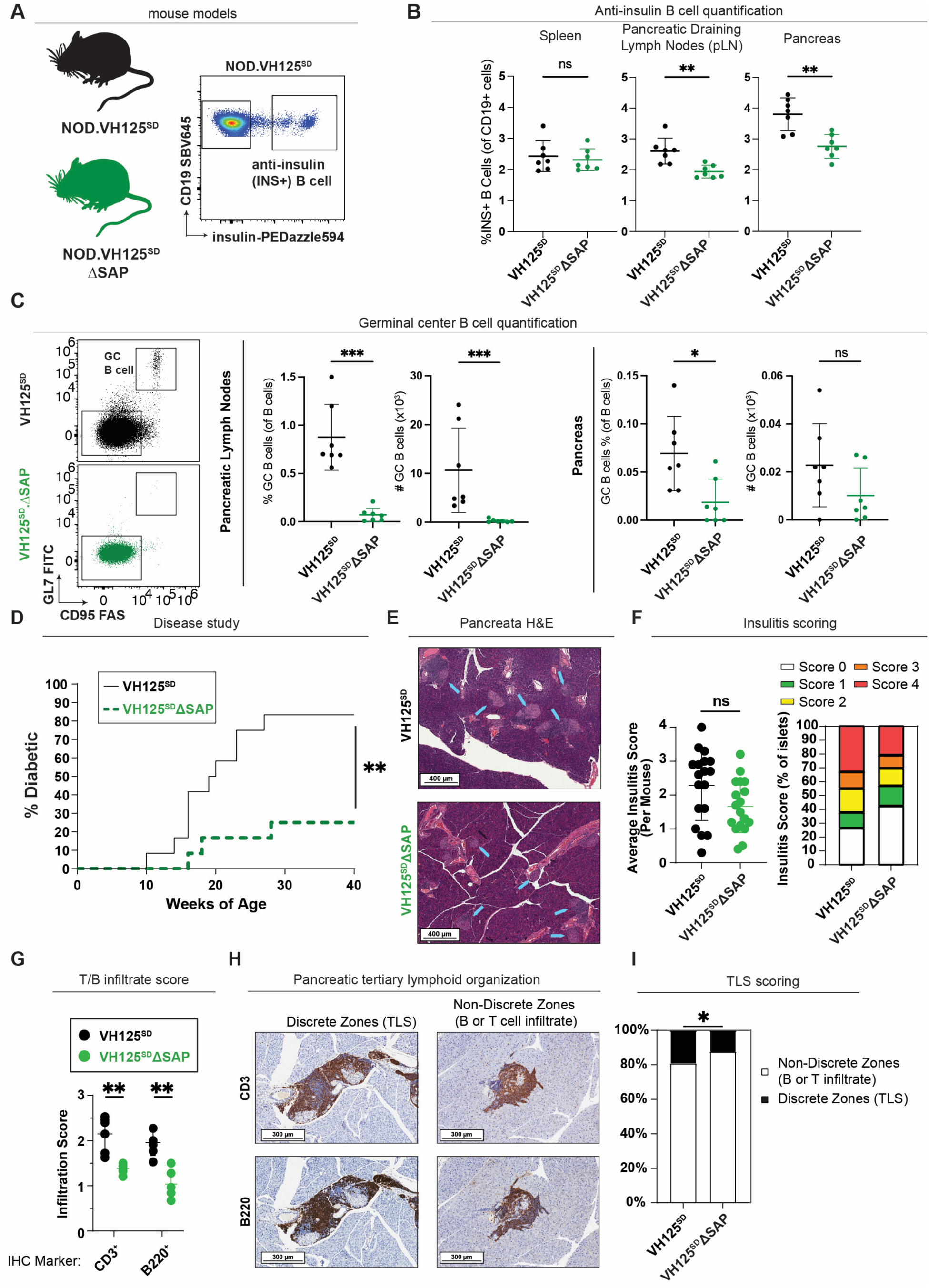
Loss of SAP reduces GC formation and T1D incidence, while insulitis persists. **(A-C)** Cells were isolated from spleen, pLNs, and pancreas as in Methods from 12-16-week-old, prediabetic mice. **(A)** VH125^SD^.NOD mice with (black) and without SAP (green). Representative flow cytometry plot shows detection of insulin-binding B cells among CD19+ live singlet lymphocytes. **(B)** Flow cytometry analysis was used to quantify % anti-insulin B cells in each organ, n ≥ 6 mice per group. **(C)** Representative flow plots from pLNs identify % GC B cells (GL7+ CD95+) among the CD19+ CD45+ live singlet lymphocyte gate (*left*), with individual mice and means plotted for pLNs (*middle*) or pancreas (*right*), n ≥ 6 mice per group. **(D)** Diabetes was monitored weekly in female VH125^SD^ΔSAP (dashed green line) and co-housed control VH125^SD^ (solid black line) mice starting at 10 weeks of age. Diabetes was diagnosed at the first of two consecutive blood glucose values ≥ 250 mg/dL. n = 12 mice/group, *p =* 0.003, log-rank test. **(E-I)** Pancreata were freshly harvested from female, prediabetic, control VH125^SD^ and VH125^SD^ΔSAP mice (8-19 weeks of age) and were formalin fixed and paraffin embedded as described in Methods. **(E)** Representative H&E-stained sections; blue arrows point to islets. **(F)** Islets were blind scored from n = 18 mice/group. *Left:* Average insulitis scores for individual mice are plotted for control (black circles) and VH125^SD^ΔSAP (green circles) mice. not significant (ns) (*p* = 0.058), Mann-Whitney *U* test. *Right:* The percentage of islets with each score is shown (0 = no insulitis, 1 = 0-25% insulitis, 2 = 25-50% insulitis, 3 = 50-75% insulitis, 4 = >75% insulitis). **(G-I)** Pancreas sections were stained via immunohistochemistry for B220^+^ or CD3^+^ expression (brown) and hematoxylin (blue) and blind scored. **(G)** Average lymphocyte infiltration score is shown for CD3^+^ and B220^+^ staining in control (black circles) and VH125^SD^ΔSAP mice (green circles). **(H)** Representative images of anti-CD3 (top) or anti-B220 (bottom) stained serial sections from control mice showing discrete B and T cell zones in tertiary lymphoid structures (TLS) or non-discrete B or T cell zones in islets. **(I)** Infiltrated islets (which excluded islets scored as peri-insulitis or no/limited insulitis) were categorized as having discrete (black bars) or non-discrete TLS (white bars) in control mice compared to VH125^SD^ΔSAP mice, n = 6 mice/group. * p ≤ 0.05, ** p ≤ 0.01, Mann-Whitney *U* test (B, C, F, G), log-rank test (D), or χ^2^ test (I).

### Loss of SAP impairs T1D incidence but not insulitis or tertiary lymphoid structure organization in insulin-skewed VH125^SD^ NOD mice

To evaluate whether loss of SAP prevented T1D development in VH125^SD^ΔSAP NOD mice, we assessed diabetes incidence. We found that VH125^SD^ΔSAP NOD mice were protected against T1D compared to VH125^SD^.NOD controls (**Figure 1D**). In alignment with other studies, in which insulitis can persist despite diabetes protection^36,37^, we observed that insulitis persisted in VH125^SD^ΔSAP NOD mice (**Figure 1E-F**), although average insulitis scores trended downward relative to controls (**Figure 1F**, p = 0.058).

To specifically evaluate T and B lymphocyte infiltration and organization, serial pancreas sections from mice which received an average insulitis severity score of 3 or 4 across the entire pancreas section (defined as in Methods) were further stained for CD3 and B220, respectively. SAP loss reduced, but did not fully abrogate, T and B cell infiltration of islets (**Figure 1G**). We further characterized T and B lymphocytic infiltrates in terms of organization, with both “non-discrete” T/B lymphocytic infiltrate and “discrete” tertiary lymphoid structures composed of organized B and T cell zones present in both SAP-sufficient and -deficient VH125^SD^.NOD mice (**Figure 1H**). The majority of T/B infiltrates had non-discrete T/B cell zones, with a small but significant decrease in the proportion of islets with discrete zones noted in ΔSAP-relative to control VH125^SD^.NOD mice (**Figure 1I**). Together, these results highlight that SAP loss protects against diabetes development even when anti-insulin B cells are present in the peripheral repertoire, despite persistent B and T lymphocyte infiltration of pancreatic islets.

### Loss of SAP impairs anti-insulin B cell proliferation, co-stimulatory molecule upregulation, and activation

Anti-insulin B cells can spontaneously adopt GC B cell phenotypes in T1D-associated organs^38^. Tfh help to B cells is compromised in the absence of SAP, even when phenotypically defined Tfh populations persist^39,40^. As shown in **(Figure 2A)**, anti-insulin and non-insulin binding B cells spontaneously adopted GC B cell phenotypes, as expected, that was subsequently lost with SAP. Given that the majority of insulin-binding B cells reside in anatomically defined EF niches^25^, we hypothesized that anti-insulin B cell activation and proliferation would be altered by loss of SAP. To eliminate the bias of differences in GC presence with/without SAP, we assessed GL7-CD95-(non-GC) insulin+ or insulin-B cells in terms of their proliferation (Ki67) or upregulation of T cell costimulatory (CD86) or activation markers (CD44). Anti-insulin B cell proliferation (% Ki67+) was increased in both pLNs (10%) and pancreas (22%), relative to either non-insulin-binding B cells (∼2-5%) in the same organs or to insulin-binding B cells in the spleen (3%) (**Figure 2B-C**). This enhanced anti-insulin B cell proliferation at the site of autoimmune attack was reduced by loss of SAP, returning it to non-insulin-binding B cell proliferation levels (∼2-5%) (**Figure 2B-C**). Anti-insulin B cells in the pancreas showed elevated expression of the T cell co-stimulatory molecule, CD86 (∼2-fold higher) relative to non-insulin-binding B cells, which was reduced by SAP loss in the pancreas, with values trending lower in pLNs, and no change in the spleen (**Figure 2D**). We observed limited differences in B cell expression of CD44 across the spleen and pancreas, with some decrease noted in the pLNs (**Figure 2E**). Overall, these data suggest that SAP supports enhanced anti-insulin B cell proliferation and CD86 upregulation, particularly in the pancreas, relative to non-insulin-binding B cells present in the same organs.

**Figure 2:**
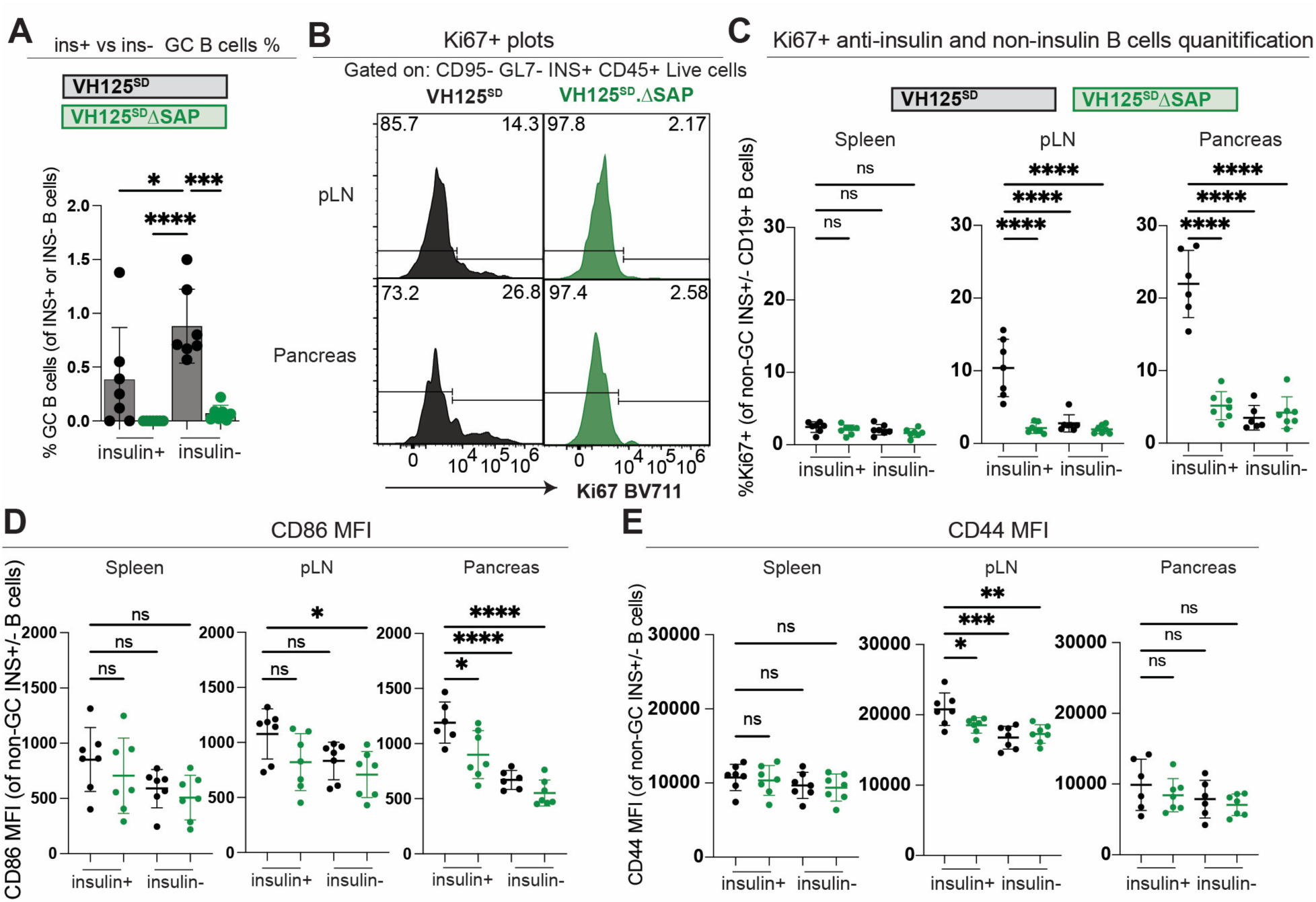
Loss of SAP impairs non-GC anti-insulin B cell proliferation, upregulation of the T cell co-stimulatory marker, CD86, and the activation marker, CD44, which is enhanced compared to non-insulin-binding B cells. Cells were isolated from spleen, pLNs, and pancreas as in Methods from 12-16-week-old, prediabetic VH125^SD^.NOD mice with (black) and without SAP (green), n=6-8 mice per group, and gated on insulin-binding B cells or non-insulin binding B cells. Data points represent individual mice. **(A)** % GC B cells (GL7+ CD95+) among insulin+ or insulin-CD19+ live singlet lymphocytes. **(B)** Representative histograms and **(C)** summaries show % Ki67+ among non-GC (Fas-GL7-), insulin-binding B cells in pLNs (*top*) and pancreas (*bottom*) from VH125^SD^ and VH125^SD^ΔSAP mice. **(D-E)** The MFI of **(D)** CD86 (T cell co-stimulatory molecule), and **(E)** CD44 (activation marker) is shown for spleen (*left*), pLNs (*middle*), and pancreas (*right*). Bars represent mean +/- standard deviation. * p ≤ 0.05, ** p ≤ 0.01, *** p ≤ 0.001, **** p ≤ 0.0001, One-Way ANOVA test with post-hoc Tukey’s multiple comparisons.

### SAP is dispensable for extrafollicular antibody responses following insulin immunization

In a model of systemic lupus erythematosus, SAP loss abolished GC responses while EF plasmablast responses remained intact^21^. Given the predominance of anti-insulin B cells in the EF niche^25^, we asked whether loss of SAP altered EF responses, with initial examination of antibody production. Immune tolerance mechanisms prevent anti-insulin B cell differentiation into antibody-secreting cells following T-dependent immunization with classic adjuvants (CFA and SRBCs), but an alternative immunization strategy that can engage TLRs overcomes this block in VH125^SD^.NOD mice to drive robust anti-insulin GC formation and anti-insulin antibody production on day 5^31,38^. Specifically, this immunization consists of insulin conjugated to *Brucella abortus* ring test antigen (insulin-BRT), which is a combination of heat-killed gram-negative bacteria coupled to insulin. The insulin-BRT immunization system thus represents a unique setting in which T-independent stimuli are present (TLR4), with insulin-BRT confirmed to drive T-independent responses in athymic mice^41,42^. However, anti-insulin GCs and IgG2a are induced by insulin-BRT immunization, suggesting a T cell help component is contributing to the immune response observed^31^. Any anti-insulin B cell responses induced by this immunization approach in the absence of SAP should arise via an EF response given the loss of GC B cells (**Figure 1C**). We therefore immunized VH125^SD^ and VH125^SD^ΔSAP mice with insulin-BRT or unconjugated BRT (BRT only) and tracked anti-insulin responses as in **Figure 3A**.

**Figure 3:**
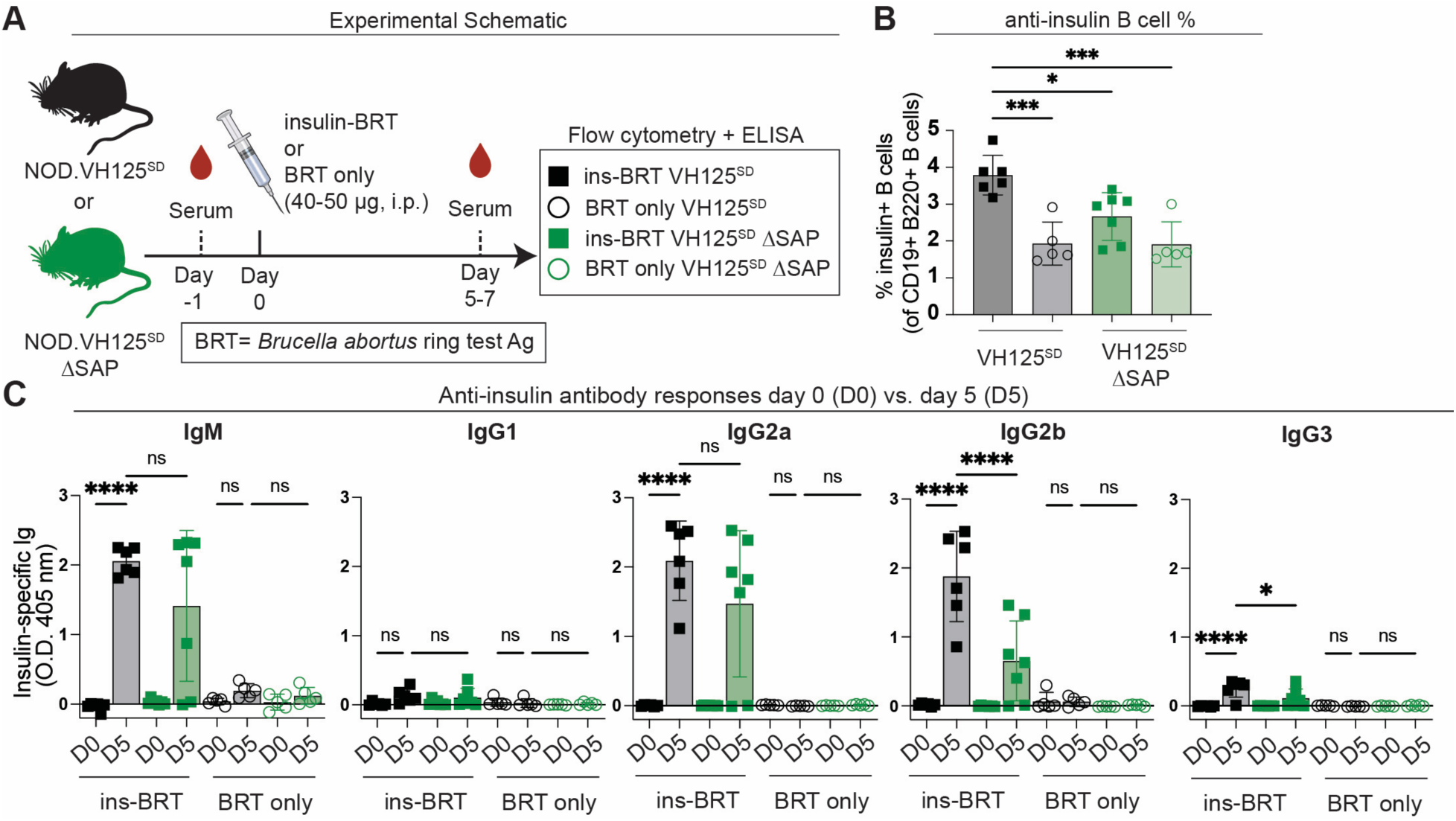
SAP is dispensable for extrafollicular antibody responses following insulin immunization. **(A)** Experimental schematic. VH125^SD^ prediabetic mice with and without SAP were immunized with insulin-*Brucella abortus* ring test antigen (ins-BRT) or BRT-only. Sera were harvested for ELISA on Day 0 (D0) or D5-7 (D5) post-immunization, with spleens collected only on D5-7 post-immunization for flow cytometry analysis. **(B)** % Insulin-binding B cells (among total B cells – CD19+ B220+ live singlet lymphocytes). **(C)** ELISA measured anti-insulin IgG1, IgG3, IgG2a/c, and IgM. Data points represent individual mice, error bars represent mean +/- standard deviation. * p < 0.05, ** p < 0.01, *** p < 0.001, **** p < 0.0001, one-Way ANOVA, followed by post-hoc Tukey’s multiple comparisons test to compare D0 vs. D5 VH125^SD^ with ins-BRT, D0 vs. D5 VH125^SD^ BRT only, D5 vs. D5 ins-BRT VH125^SD^ vs. VH125^SD^ ΔSAP, and D5 vs. D5 BRT only VH125^SD^ vs. VH125^SD^ ΔSAP.

Anti-insulin B cells expanded in the spleen following insulin-BRT, but not BRT-only immunization, with somewhat lower expansion in SAP-deficient mice (**Figure 3B)**. Insulin-BRT immunization elicited anti-insulin antibodies of the IgM, IgG2a, and IgG2b isotypes in the sera of all VH125^SD^ and most VH125^SD^ΔSAP mice, with the BRT-only negative control failing to drive an antibody response (**Figure 3C**). Of note, TNP-ficoll (T-independent) immunization drives IgG1 and IgG3 responses in the absence of SAP^43^, but these classic T-independent isotypes were not induced by insulin-BRT immunization (**Figure 3C**).

### SAP supports anti-insulin GC and atypical B cell expansion following insulin immunization

To directly evaluate how anti-insulin B cell responses were impacted by SAP loss, we utilized a high-dimensional spectral flow cytometry panel followed by minimally supervised analysis to assess B cell phenotypic changes following insulin-BRT or control BRT-only immunization. As shown in **Supplemental Figure 2A**, we gated on insulin+ B cells and identified n = 22 distinct clusters using Phenograph^34^, with marker expression t-SNEs shown in **Supplemental Figure 2B**. We collapsed these clusters into n = 8 populations: GC, follicular, CD138+ TACI+ (antibody-secreting cells), activated, atypical, marginal zone, CD73+ (memory-like), or CD80+PDL2+ (memory-like)^44^ (**Figure 4A)**, with heatmap expression of each marker shown for each population (**Figure 4B**). Differential abundance analysis using diffcyt was used to identify population changes across groups, defined as |log2FC| > 1 and p < 0.01^35^. Following insulin-BRT immunization in VH125^SD^ mice, activated, GC, atypical, and CD138+ TACI+ B cells expanded within anti-insulin B cells (**Figure 4C).** Loss of SAP reduced GC and atypical B cell populations within anti-insulin B cells following insulin-BRT immunization (**Figure 4C**). These changes were also identified in the original Phenograph clusters (**Supplemental Figure 2C**). However, no differences were found in antibody-secreting cells or memory-like CD80+ PDL2+ populations with SAP loss using either diffcyt or manual gating analysis (**Figure 4C-G**). Immunization with insulin-BRT also drove induction of CD80^+^ PDL2^-^and CD80^-^PDL2^+^ memory-like populations (**Supplemental Figure 3**). Together, these results show that insulin-BRT immunization drives anti-insulin IgG and memory-like B cell formation independently of SAP, whereas anti-insulin GC and atypical B cell responses are reduced by SAP loss.

**Figure 4:**
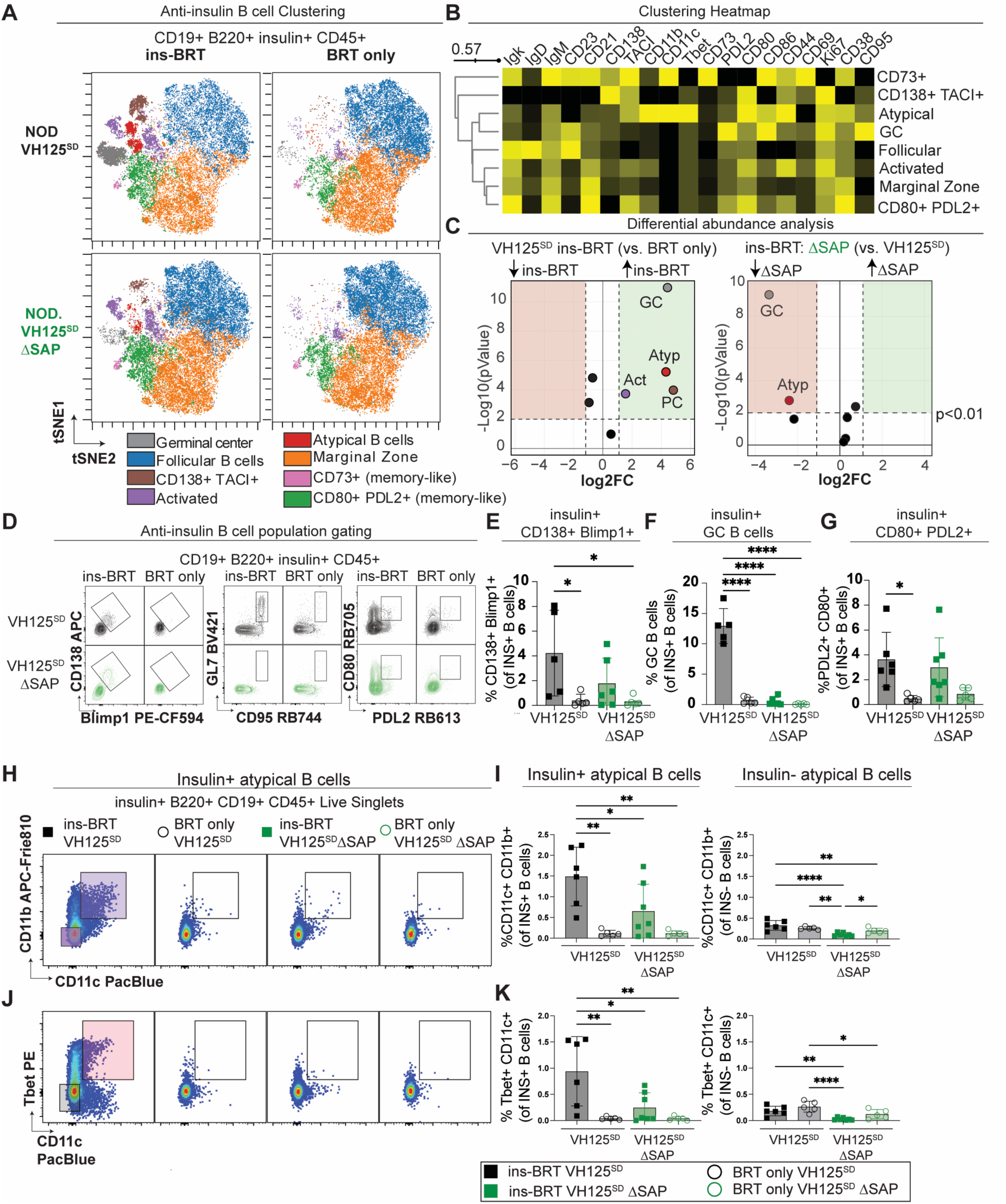
SAP supports anti-insulin GC and atypical B cell formation but is dispensable for anti-insulin memory-like B cell and antibody-secreting cell formation. **(A-C)** Anti-insulin B cells (CD45+ B220+ CD19+ insulin+ live singlet lymphocytes) underwent minimally supervised analysis via Phenograph, as in **Supplemental** Figure 2A, to identify n = 22 clusters which were manually collapsed into the n = 8 populations shown on (**A)** t-SNE plots for VH125^SD^ and VH125^SD^ ΔSAP mice following ins-BRT and BRT-only immunization. **(B)** Heatmap showing representative marker expression for each cluster in *(A)*. **(C)** Differential abundance analysis for each cluster via diffcyt comparing either ins-BRT vs BRT-only immunization in VH125^SD^ SAP-sufficient mice (*left*), or ins-BRT immunization comparing SAP-sufficient and -deficient VH125^SD^ mice (*right*). **(D)** Flow cytometry gating schemes identify antibody-secreting cells (CD138+ Blimp1+, *left*), GC B cells (GL7+ CD95+, *middle*), and memory-like B cells (CD80+ PDL2+, *right*) among insulin-binding B cells (CD19+ B220+ CD45+ Insulin+ live singlet lymphocytes). **(E-G)** The proportion of each population among insulin+ B cells for **(E)** antibody-secreting cells, **(F)** GC B cells, or **(G)** memory-like B cells is plotted for individual mice. **(H-K)** Atypical B cell subsets were defined as either **(H-I)** CD11b+ CD11c+ or **(J-K)** CD11c+ T-bet+ among insulin+ or insulin-B cells (CD45+ B220+ CD19+ live singlet lymphocytes). **(H)** Representative flow plots of CD11b+ CD11c+ subset, or (**J)** T-bet+ CD11c+ subset. **(I)** Quantification of % CD11c+ CD11b+ or **(K)** % T-bet+ CD11C+ amongst insulin+ B cells (*left*) or insulin-B cells (*right*). n = 5-7 individual mice per group, bars represent mean +/- standard deviation. * p < 0.05, ** p < 0.01, *** p<0.001, **** p<0.0001, one-way ANOVA with post-hoc Tukey’s multiple comparisons test.

Given the reduction in atypical B cell induction by this immunization in the absence of SAP, we separately evaluated CD11b+ CD11c+ and CD11c+ T-bet+ subsets, as in **Figure 4H,J**. Insulin-BRT immunization provoked expansion of CD11b^+^ CD11c^+^ (**Figure 4I**) and T-bet^+^ CD11c^+^ (**Figure 4K**) populations compared to BRT-only immunization, which was reduced by loss of SAP (**Figure 4I,K)**. Expansion of these atypical B cell populations was not observed in non-insulin-binding B cells **(Figure 4I,K)**. These results show that SAP supports expansion of both atypical B cell populations following insulin-BRT immunization.

### Anti-insulin atypical B cells upregulate activation, T cell co-stimulatory, and proliferation markers, and undergo IgG2a switching following insulin-BRT immunization

To further evaluate whether the atypical B cells that expanded following insulin-BRT immunization appeared poised to act as antigen-presenting cells to T cells, we next measured CD80 and CD86 expression, along with activation markers CD44 and CD69, and the proliferative marker, Ki67, relative to non-atypical B cells. Specifically, we evaluated 1) anti-insulin CD11c^+^ CD11b^+^, 2) anti-insulin T-bet^+^ CD11c^+^, 3) anti-insulin T-bet^-^CD11c^-^, 4) anti-insulin CD11c^-^CD11b^-^, and 5) non-insulin-binding B cell populations following insulin-BRT immunization of VH125^SD^ mice, as in **Figure 5A**. Anti-insulin CD11c+ T-bet+ atypical B cells expressed higher levels of CD86, CD80, CD44, and CD69, with trending increases in Ki67 compared to the T-bet^-^CD11c^-^insulin-binding B cells or the non-insulin-binding B cells (**Figure 5B-F**). Similar upregulated expression of CD86, CD80 (co-stimulatory) and CD44 (activation) was seen in CD11b+ CD11c+ anti-insulin B cells compared to CD11b-CD11c-anti-insulin B cells, except for a trending increase in CD69 (activation) expression (**Figure 5B-F**). Lack of CD21 and CD23 expression is also a hallmark feature of age-associated/atypical B cells^45,46^. Consistent with this, we found that T-bet+ CD11c+ atypical B cells were mostly (∼70%) CD21-CD23-, which decreased to ∼50% in CD11b+ CD11c+ atypical anti-insulin B cells (**Figure 5G).**

**Figure 5:**
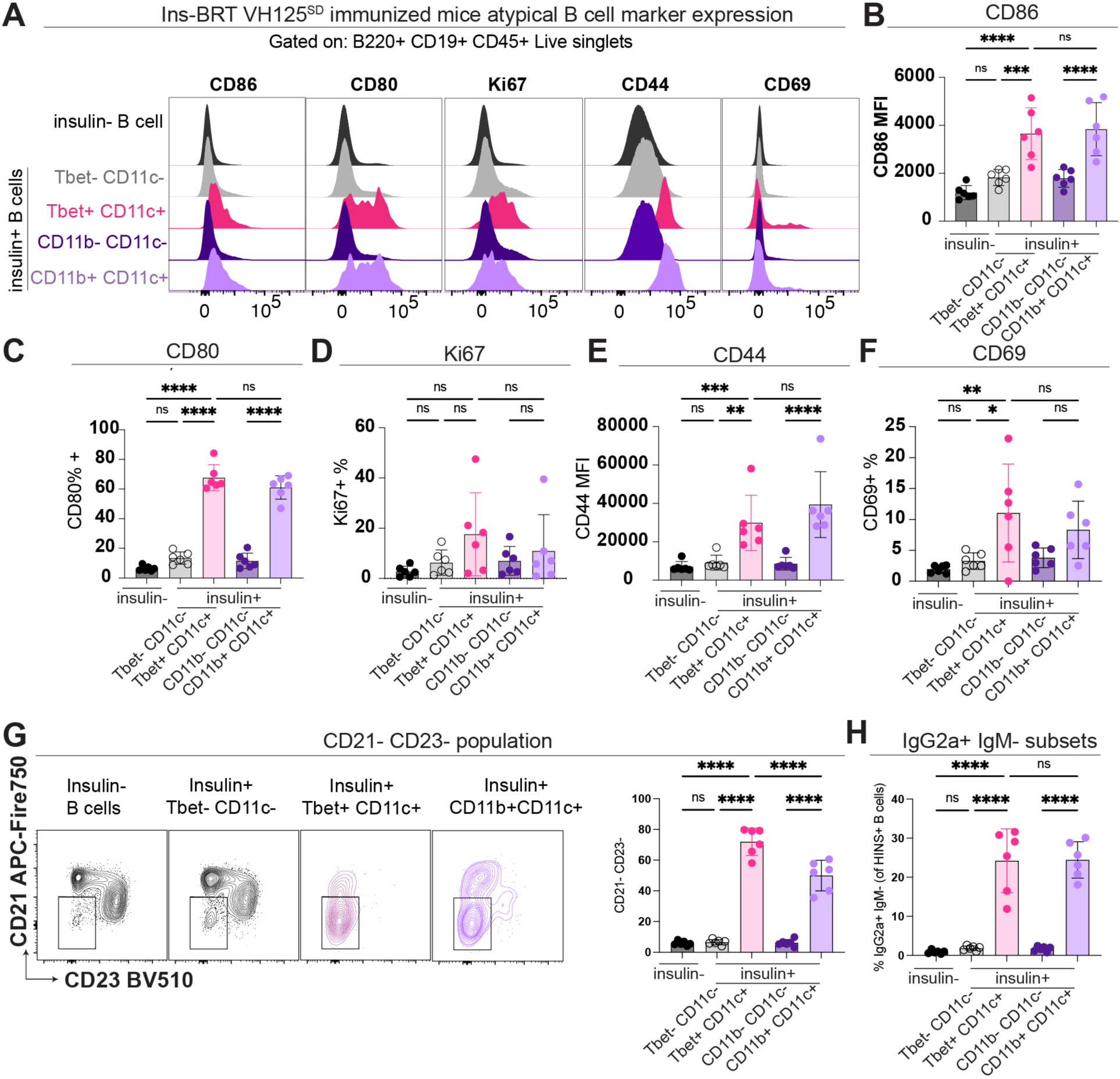
Atypical insulin-binding B cells show enhanced activation, proliferation, and upregulation of T cell costimulatory molecules following insulin-BRT immunization. VH125^SD^ prediabetic mice were immunized with insulin-*Brucella abortus* ring test antigen (ins-BRT), and flow cytometry analysis of splenocytes was performed 5-7 days after immunization. B cells were further gated on insulin-B cells (black), T-bet-CD11c-insulin+ (grey), T-bet+ CD11c+ insulin+ (pink), CD11b-CD11c-insulin+ (dark purple), and CD11b+ CD11c+ insulin+ (light purple). Data points represent individual mice. **(A)** Histogram plots show expression of CD86 (T-cell costimulatory molecule), CD80 (activation), Ki67 (proliferation), CD44 (activation), and CD69 (activation). **(B-F)** Individual mice are plotted for **(B)** CD86 MFI, **(C)** CD80 MFI, **(D)** % Ki67+, **(E)** CD44 MFI, and **(F)** % CD69+. **(G)** % CD21-CD23-within each of these subsets is shown in representative plots (*left*) and summarized (*right*). **(H)** % IgG2a+ B cells among each B cell subset shown. n = 5-7 mice per group, bars represent mean +/- standard deviation. * p < 0.05, ** p < 0.01, *** p < 0.001, **** p < 0.0001, one-way ANOVA with post-hoc Tukey’s multiple comparisons test.

IgG2a can arise in response to T-independent immunization in a T-bet-dependent manner^47^, a transcription factor upregulated in some atypical B cells^48^. Roughly 25% of anti-insulin T-bet^+^ CD11c^+^ and CD11b+ CD11c+ B cells switched to the IgG2a isotype, with non-atypical B cell subsets showing little to no IgG2a switching **(Figure 5H)**. Overall, insulin-BRT immunization drives the formation of anti-insulin atypical B cells, which upregulate T cell co-stimulatory molecules that could support their function as antigen-presenting cells. This was associated with heightened atypical B cell activation, proliferation, and IgG2a class switching.

### T follicular helper (Tfh) and T peripheral helper (Tph) phenotype cells form independently of SAP in anti-insulin-skewed VH125^SD^.NOD mice

SAP is essential for long-lived cognate T-B interactions, and, when blocked, abrogates productive GC-derived antibody responses^49–51^. These responses depend on SAP expression in T cells, but not B cells^39,52^. The role of SAP in autoimmune disease is nuanced, as SAP-deficiency led to near complete loss of Tfh cells and GC B cells in the K/BxN model of autoimmune arthritis, yet Tfh cells (CXCR5^hi^ PD-1^hi^ BCL6^+^ ICOS^hi^ CD44^hi^) persisted in the NOD model of T1D despite GC B cell collapse in ΔSAP NOD mice relative to NOD controls^29^. The persistence of CD4+ T cells with a Tfh phenotype in some settings of SAP-deficiency can in part be explained by the finding that CD4+ T cells can still upregulate the Tfh markers, CXCR5 (which supports B cell follicle homing) and BCL6 (a transcriptional repressor that supports Tfh maturation) even when SAP is lost ^51,53^.

As shown in **Figure 6A**, a phenotypically defined Tfh population (Foxp3-CXCR5^hi^ PD-1^hi^) persisted in the pLNs and pancreas of VH125^SD^ΔSAP mice. BCL6 is an essential transcriptional repressor for Tfh cell differentiation ^54,55^. Regardless of SAP, BCL6 expression was detected in a higher frequency of Tfh cells compared to non-Tfh cells in pLNs and pancreas (**Figure 6B**), consistent with BCL6 functioning upstream of SAP to support Tfh maturation^56^. T peripheral helper cells (Tph, ICOS^hi^ PD-1^hi^ CXCR5^-^) are found in inflamed tissues in other autoimmune contexts^57^, but their dependence on SAP is unclear. We found no differences in the frequency of Tph cells in either pLNs or pancreas when SAP was lost (**Supplement Figure 5A-B**). Together, these results show that phenotypically defined Tfh and Tph cells persisted despite loss of GC B cells with SAP-deficiency in the VH125^SD^.NOD model.

**Figure 6:**
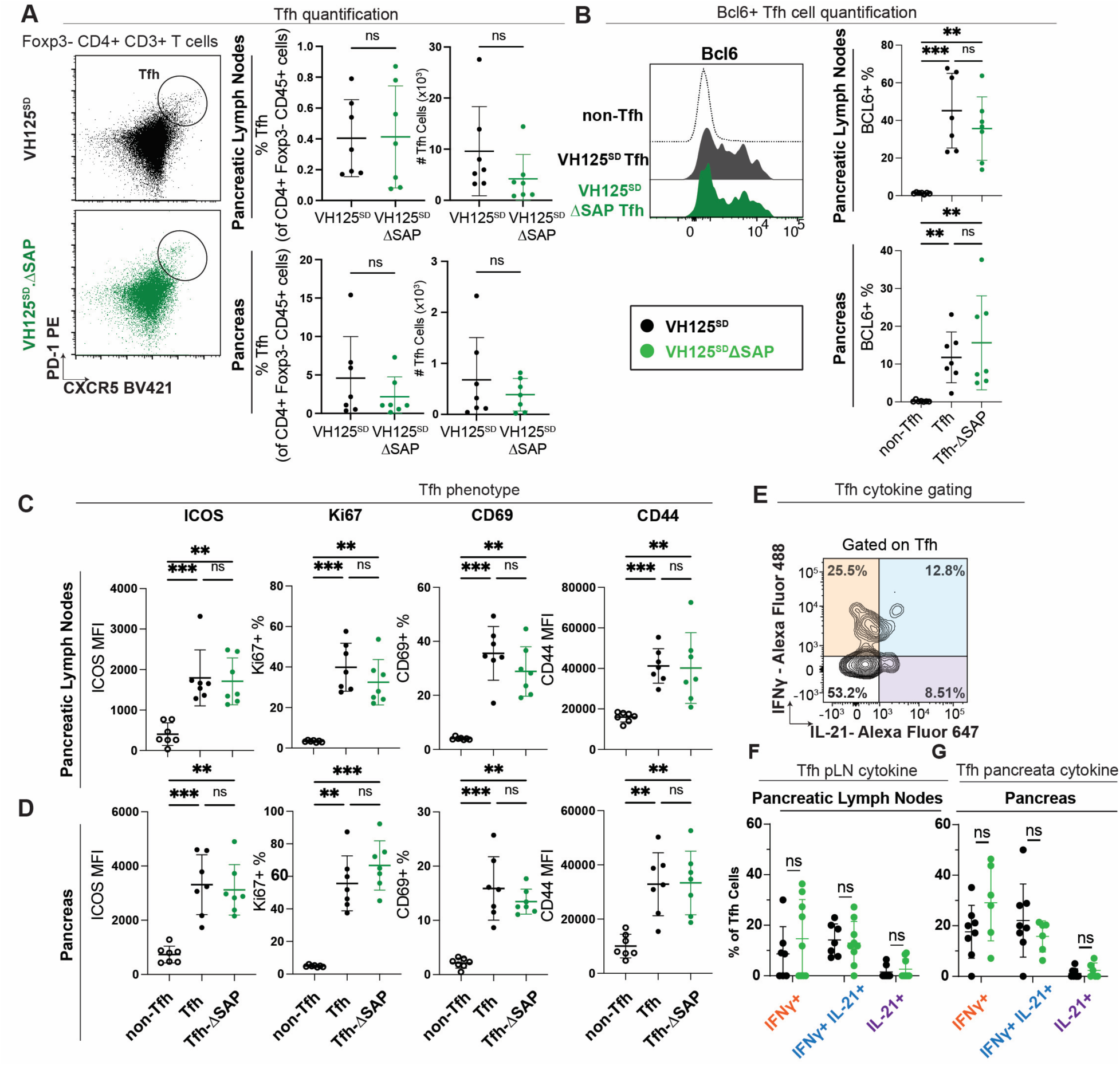
Loss of SAP does not impact Tfh formation or cytokine production. Pancreas and pLNs were isolated from 12-16-week-old, prediabetic VH125^SD^ mice with and without SAP and underwent flow cytometry analysis. Individual data points represent individual mice, n=6-8 mice per group. **(A)** Representative flow cytometry plot identifies % Tfh cells (CXCR5hi PD-1hi) among CD4+ Foxp3-CD45+ live lymphocytes (*left*), with summary data showing Tfh frequency and number within pLNs (*top right*) and pancreas (*bottom right*). **(B)** Representative histograms show BCL6 expression within non-Tfh (dotted), VH125^SD^ Tfh (grey) and VH125ΔSAP Tfh mice (green) pLNs (*left*). % BCL6+ cells among Tfh-gated cells in pLNs (*top right*) and pancreas (*bottom right*). **(C-D)** ICOS MFI, % Ki67+, % CD69+, and CD44 MFI for non-Tfh, Tfh, and Tfh-ΔSAP groups within **(C)** pLNs and **(D)** pancreas. **(E)** Representative flow plot of Tfh IFN-γ and IL-21 production, identified as in Methods. **(F-G)** Frequency of IFN-γ+, IL-21+ IFN-γ+, and IL-21+ cells amongst Tfh in **(F)** pLNs and **(G)** pancreas. For all analyses, bars represent mean +/- standard deviation, * p ≤ 0.05, ** p ≤ 0.01, *** p ≤ 0.001, Mann Whitney U test (A) or Kruskal-Wallis test post hoc Dunn’s test (B-G).

### Loss of SAP does not reduce Tfh or Tph upregulation of activation markers, proliferation, or pro-inflammatory cytokine skewing in VH125^SD^.NOD mice

SAP-deficient Tfh cells can still upregulate CXCR5 and BCL6^51,53^, though some studies show ICOS upregulation is compromised^39^. However, downstream Tfh functions are impaired by loss of SAP, including the prolonged Tfh:B cell contact requited to form GC B cells, drive GC Tfh maturation, and support plasma cell differentiation ^14,51,58^. Given that we observed Tfh cell retention despite reduced diabetes incidence in VH125^SD^.SAP^-/-^NOD mice vs. SAP-sufficient controls, we next tested whether Tfh cellular function and/or phenotype were altered by assessing markers of Tfh activation (ICOS, CD44, CD69), recent proliferation (Ki67), chemokine receptor expression (CXCR3, CCR6), and cytokine production (IL-21, IFN-γ). No changes in ICOS, Ki67, CD69, and CD44 were found in pLNs or pancreas isolated from VH125^SD^ΔSAP vs. VH125^SD^ mice, with all these markers upregulated in Tfh vs. non-Tfh populations (**Figure 6C-D**). Tfh cells preferentially skewed towards CXCR3^+^ CCR6^-^(Tfh1-like) and CXCR3^-^CCR6^+^ (Tfh17-like) populations compared to non-Tfh counterparts (**Supplemental Figure 4A-B**), which was not altered by loss of SAP (**Supplemental Figure 4C**).

Tfh and Tph populations can express the pro-inflammatory cytokines, IFN-γ and/or IL-21 ^57,59,60^. The frequencies of IFN-ψ+ and IL-21+ IFN-γ+ populations in the pLNs and pancreas ranged from <1-5% within total CD4 vs. ∼20-30% within Tph and ∼10% within Tfh, highlighting enhanced skewing of Tph and Tfh towards these pro-inflammatory cytokines (**Figure 6** and **Supplemental Figure 5**). SAP loss in the VH125^SD^ model did not alter the frequencies of IFN-γ+, IL-21+, or IL-21+ IFN-γ+ populations within 1) total CD4+ T cells (**Supplemental Figure 5C-D**), 2) Tfh (**Figure 6E-G**), or 3) Tph (**Supplemental Figure 5E-F**) subsets in the pLNs or pancreas, with the exception that Tph in the pancreas showed a small, but statistically significant % IFN-γ+ increase and % IFN-γ+ IL-21+ decrease (**Supplemental Figure 5E-F**). Together, these results suggest that these facets of Tfh programming are not altered by loss of SAP in VH125^SD^ mice, implicating other mechanisms of diabetes protection in this setting.

### SAP loss reduces CD44 and CCR6 expression on T-B doublets formed *in vivo* in VH125^SD^.NOD mice

Given the persistence of Tfh and Tph populations in VH125^SD^ΔSAP mice (**Figure 6** and **Supplemental Figure 5**), we next tested whether B-CD4 T cell conjugate formation *in vivo* was altered by loss of SAP, with B-CD4 T cell conjugates identified amongst insulin+ (INS+) or insulin-(INS-) cells as in **Figure 7A**. We evaluated B-CD4 T cell conjugates in pLNs and spleen, but not pancreas, as the more extensive protocol required to isolate pancreas single-cell suspensions limited conjugate detection in this tissue (data not shown). B-CD4 T cell conjugates were larger in size compared to CD19+ CD4-and CD19-CD4+ single positive populations, as expected (**Figure 7B**). We focused on non-GC B cells to eliminate bias between VH125^SD^ mice (GC B cells present) and VH125^SD^ΔSAP mice (GC B cells absent). A 2-3-fold increased frequency of INS+ relative to INS-B-T conjugates was observed in both pLNs (∼3% vs. ∼1%) and spleen (∼5% vs. ∼2%), which was not reduced by SAP loss (**Figure 7C).** We next tested whether B-T conjugates exhibited altered activation (CD44), upregulation of co-stimulatory molecules (ICOS, CD86), pro-inflammatory potential (CCR6), or proliferation (Ki67) in pLNs or spleen. CD44 and CCR6 expression was elevated on T-B conjugates in the pLNs relative to spleen and was significantly decreased on INS+ B-T conjugates by loss of SAP in the pLNs (**Figure 7D-E**). No differences were observed in T-B conjugate expression of ICOS, CD86, or in the frequency of Ki67+ cells (**Supplemental Figure 6A-C**). GC B-T cell conjugates were reduced in VH125^SD^ΔSAP vs. VH125^SD^ mice, as expected (**Figure 7F**). Together, these results suggest that loss of SAP impacts some, but not all, of the activation and proinflammatory markers evaluated here.

**Figure 7:**
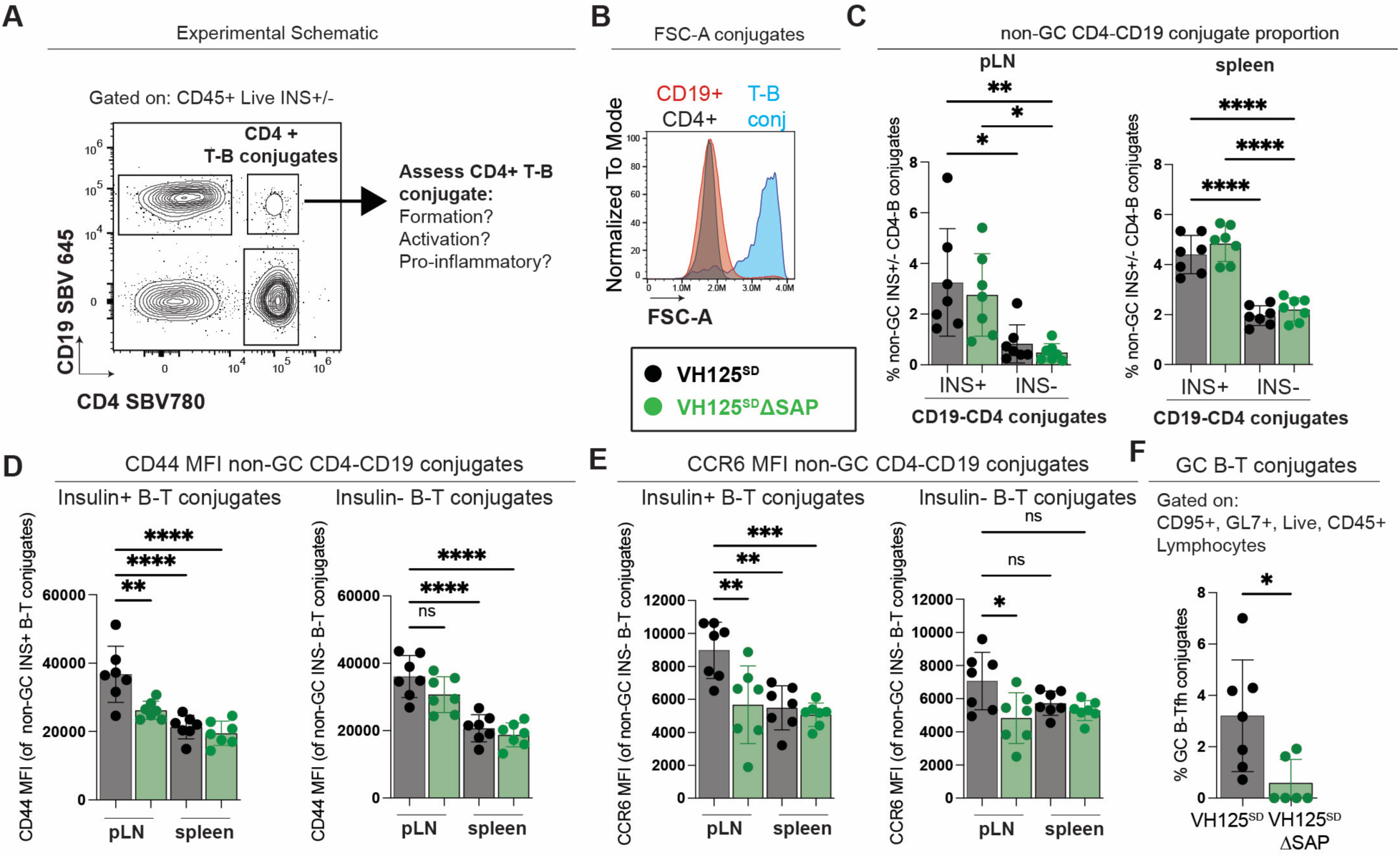
Anti-insulin B cells show an increased propensity to form B-CD4+ T cell conjugates *in vivo* relative to non-insulin-binding B cells, with SAP-deficiency causing reduced CD44 and CCR6 expression. Spleen and pLNs were isolated from 12-16-week-old, prediabetic VH125^SD^ and VH125^SD^ ΔSAP mice. **(A)** Experimental schematic. CD4+ T-B conjugates were identified by gating on CD45+ live lymphocytes, and then gated on insulin+, or insulin-, followed by CD19+ CD4+ to obtain CD4+ T-B conjugates. **(B)** FSC-A (size) of cells in either CD4+ T-B conjugates, CD19+ CD4-cells, or CD4+ CD19-cells. **(C)** % CD4-CD19 conjugates is plotted among insulin+ and insulin-non-GC (GL7-CD95-) B cells in pLNs and spleen. **(D)** CD44 MFI (activation marker) and **(E)** CCR6 MFI (pro-inflammatory marker) is shown among insulin+ and insulin-B-T conjugates in PLNs and spleen. **(F)** GC B-Tfh conjugates were identified in the pLNs by gating on CD95+ GL7+ CD4+ CD19+ CD45+ live lymphocytes. For all analyses, data points represent n=6-8 mice per group, bars represent mean +/-standard deviation, * p ≤ 0.05, ** p ≤ 0.01, *** p ≤ 0.001, **** p ≤ 0.0001, one-way ANOVA followed by Tukey’s post-hoc multiple comparisons test (C-E), or Mann-Whitney U test (F).

### Loss of SAP reduces islet-reactive, progenitor exhausted-like CD8+ T cells in insulin-skewed VH125^SD^.NOD mice

CD4 T-B interactions are thought to support the downstream diabetogenic CD8+ T cell programming that enables their attack on beta cells^61^. Given the reduction in diabetes incidence in VH125^SD^ΔSAP vs. VH125^SD^ mice (**Figure 1A**), we examined islet-reactive CD8+ T cells for differences in effector programming in the pLNs and pancreas using the flow cytometry gating schemes shown in **Figure 8A, B, and D**. T cells reactive to either islet-specific glucose-6-phosphatase catalytic subunit-related protein (IGRP, mimotope NRP-V7) or insulin IA^g7^ tetramers were observed in the pLNs and pancreas of all mice, regardless of the presence/absence of SAP (**Figure 8C, E**). During states of chronic antigen exposure and in highly inflamed tissues, CD8+ T cells can adopt a semi-exhausted stem-cell-like state known as T progenitor exhausted (Tpex) and these cells can further differentiate into CD8+ effectors^62^. Tpex cells can be found within the pLNs and pancreas and are implicated in the development of islet-reactive cytotoxic CD8 T cells that contribute to T1D in NOD mice^63^. We therefore examined this population and found that insulin and IGRP-reactive Tpex cells were decreased by loss of SAP in the pancreas (**Figure 8E**), but not pLNs (**Figure 8C**) of VH125^SD^ mice. Both IGRP and insulin-reactive effector-like CD8+ T cells trended down with loss of SAP in the pancreas, with no changes observed in pLNs (**Figure 8C, E**).

**Figure 8:**
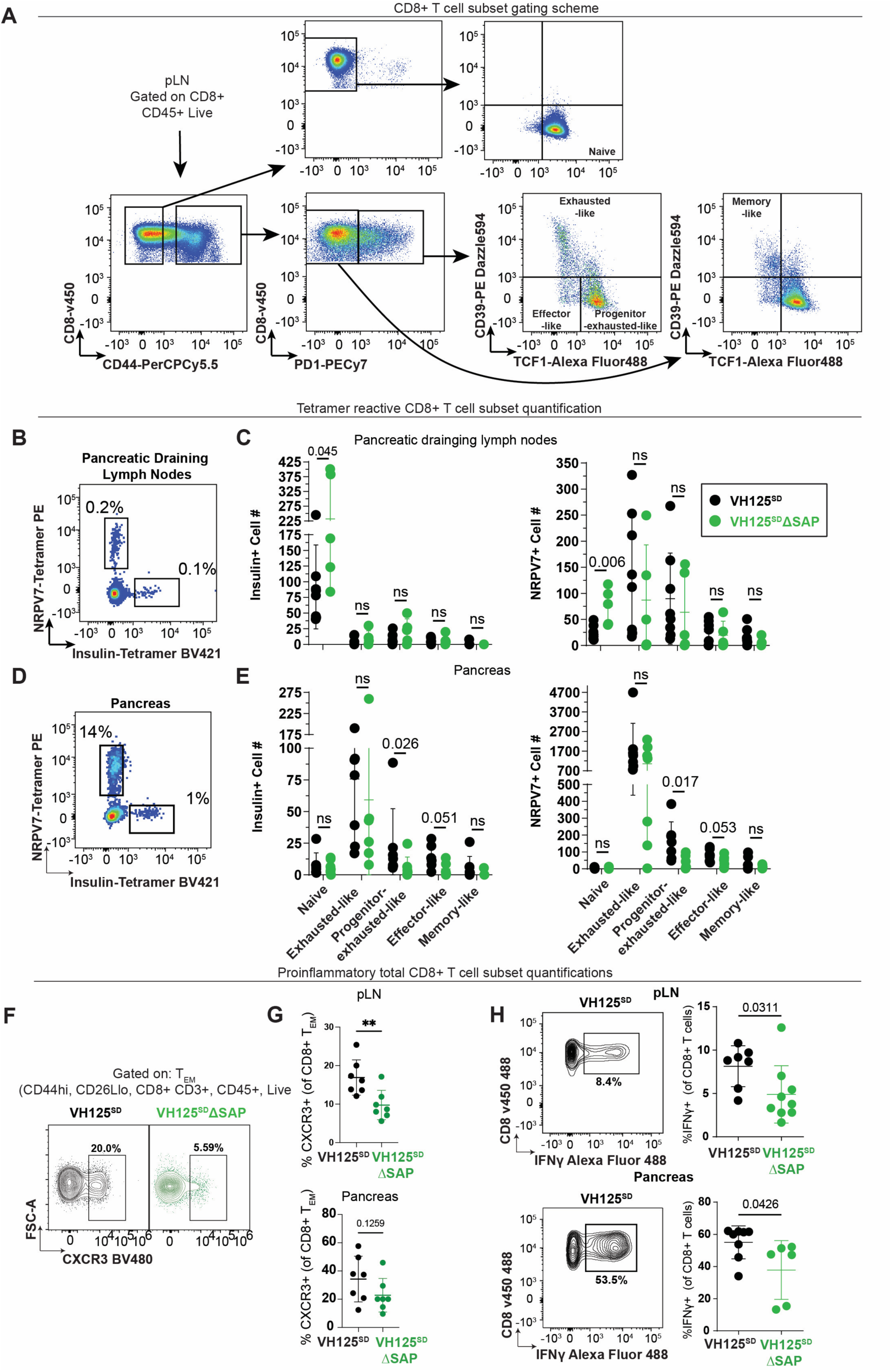
Loss of SAP reduces islet-reactive progenitor exhausted-like and IFN-γ+ pro-inflammatory CD8 T cell populations in VH125^SD^ NOD mice. Pancreas and pLNs were freshly harvested from 11-13 week-old, pre-diabetic VH125^SD^ΔSAP (SAP-deficient) and co-housed control VH125^SD^ (SAP-sufficient) mice. Cells were isolated and stained for flow cytometry analysis as in Methods. **(A)** Representative flow cytometry plots from pLNs show gating schemes used to identify CD8 populations analyzed in subsequent panels. **(B, D)** Representative flow plots of NRPV7 tetramer^+^ (IGRP-mimotope) and insulin tetramer^+^ CD8+ T cells from **(B)** pLNs and **(D)** pancreas from control (black) and VH125^SD^ΔSAP (green) mice. **(C, E)** The number of insulin reactive (*left*) and NRPV7 reactive (*right*) CD8^+^ T cell subsets in **(C)** pLNs and **(E)** pancreas. **(F)** Representative flow pots identify % CXCR3+ cells among CD8+ T effector memory (T_EM_), quantified in **(G)** for pLNs (*top*) and pancreas (*bottom*). **(H)** Representative flow plots of IFN-γ expression amongst CD45+, live, CD8+, T cells in VH125^SD^ mice (*Left*). % IFN-γ+ CD8+ T cells for individual VH125^SD^ (black) or VH125^SD^ΔSAP (green) mice in pLNs (*top*) and pancreas (*bottom*). n ≥ 5 mice individual mice plotted per group, n ≥ 3 experiments, bars represent means. * p < 0.05, ** p < 0.01, or p value listed, Mann-Whitney *U* test.

CXCR3 mediates CD8^+^ T cell homing to inflamed tissues, and subsets of CD8+ T cells such as T effector memory (T_EM_, CD44^+^ CD62L^-^) can assist in islet destruction^64^. CD8+ T cells can express IFN-γ, which drives beta cell stress and immune cell recruitment to islets, contributing to beta cell destruction^65,66^. We evaluated the total CD8^+^ T cells for changes in (1) CXCR3, gated as in **Figure 8F**, and (2) IFN-γ, gated as in **Figure 8H**. The frequency of CXCR3+ in T_EM_ CD8+ T cells was reduced in pLNs, but not pancreas, by SAP loss (**Figure 8F-G**). Additionally, the proportion of IFN-γ+ CD8+ T cells was lower in both the pLNs and the pancreas in VH125^SD^ΔSAP mice compared to SAP-sufficient controls (**Figure 8H**). These data show that SAP loss reduces islet-reactive CD8+ Tpex cells and reduces polyclonal, proinflammatory CD8+ T cell populations identified by CXCR3 and IFN-γ expression.

## Discussion

In this study, we show that despite preformed anti-insulin B cells (via an IgH transgene), loss of SAP led to a significant reduction in T1D development. Reduced diabetes occurred despite the preservation of a phenotypically defined Tfh population. However, signs of compromised T-B communication were evident with loss of SAP, as anti-insulin B cell activation, proliferation, and upregulation of T cell costimulatory molecules were diminished. Anti-insulin, atypical memory-like B cells and IgG2a-switched antibody-secreting cells expanded following an immunization approach that drove both EF and GC anti-insulin B cell expansion, with SAP loss reducing or abolishing both populations. These anti-insulin atypical B cells upregulated CD80 and CD86, suggesting they may be poised for antigen presentation. Ultimately, SAP loss was associated with reduced CD8+ T cell CXCR3+ and IFN-γ production and diminished islet-reactive CD8 Tpex cells in the pancreata and pLNs, which tracked with the diabetes protection observed.

SAP deficiency prevents the onset of lupus and rheumatoid arthritis-like disease in mice, which depend on pathologic autoantibody formation^29,67–69^, but is dispensable for the T cell-mediated autoimmune disease, experimental autoimmune encephalitis (EAE)^70^. The role of SAP in T1D is a bit more complex. Our prior study showed that 50% of NOD.ΔSAP mice still developed diabetes^29^, whereas only 25% of NOD.VH125^SD^ΔSAP mice developed diabetes here, despite the increased diabetes penetrance and acceleration noted in the VH125^SD^.NOD model ^27^. This is consistent with a larger dependence on SAP in this anti-insulin B cell-centric model, in which T1D immunopathology may require a slightly different constellation of events relative to WT.NOD mice to culminate in downstream beta cell destruction. In line with this concept, organized TLS were found in ∼50% of WT.NOD mice^29^, but only ∼20% of VH125^SD^.NOD.

SAP is dispensable for autoreactive EF antibody responses, both in our data and prior studies^21^, but SAP’s role in atypical B cell expansion was not previously studied. Our data showed that BCR/TLR co-ligation (via insulin-BRT immunization) induced anti-insulin T-bet^+^ CD11c^+^ (atypical) and GC B cells, which was reduced in the absence of SAP, whereas memory-like anti-insulin B cells and anti-insulin IgG2a still formed in response to insulin-BRT immunization. These data highlight SAP in supporting atypical B cell formation in T1D, and perhaps other autoimmune contexts. The robust induction of GC B cells within 5 days of insulin-BRT immunization could be explained by recruitment of responding cells from a pre-existing memory B cell pool. In line with this, our prior study showed that anti-insulin B cells skewed towards a CD80^+^ PDL2^+^, memory-like B cell phenotype^25^. However, given the ambiguity surrounding memory B cell phenotypic markers vs. true memory B cell fate, future studies will be required to formally evaluate whether this CD80^+^ PDL2^+^, memory-like B cell population shows other features of memory B cell programming, or if the anti-insulin memory pool is contributing to the insulin-BRT GC response.

Anti-insulin 8F10 CD4 T cells are a pathogenic, low-affinity clone observed in murine and human T1D^37^. Co-transfer of anti-insulin VH125^SD^ B cells + 8F10 T cells enhances diabetes in RAG1^-/-^.NOD recipients relative to 8F10 T cell transfer alone, highlighting the importance of these cognate B-T interactions in driving islet attack^71^. CD80/CD86 upregulation by anti-insulin atypical (relative to non-atypical) B cells suggests they may be poised for antigen-presentation. However, we did not directly evaluate their antigen-presentation capacity in this study, due to the low number of atypical B cells detected in VH125^SD^.NOD mice. B cell cross presentation of antigen via MHC class I to CD8 T cells directly has also be associated with T1D pathology^72^. Future studies will be required to evaluate theses additional routes of anti-insulin B-T cell activation, and their reliance on SAP, in driving T1D pathology.

In T1D, pro-inflammatory cytokines like IFN-γ can contribute to beta cell destruction to support cytotoxic CD8 function, suppress regulatory T cells, or contribute to beta cell destruction directly^73^. Loss of SAP led to a lower average frequency of IFN-γ-producing CD8+ T cells. Tpex are another proinflammatory population of CD8+ T cells that can produce islet-reactive effectors that traffic to the pancreas^63^. We found that SAP supports islet-reactive Tpex-like cells in the pancreas of VH125^SD^.NOD mice, with less Tpex skewing noted in the pLNs. Evidence points to roles for SAP outside of regulating SLAM-family receptor signaling, via regulation of PD1 in CD8+ T cells^74^. Tpex CD8+ T cells require PD-1 signaling to stabilize TCF1+, preventing them from differentiating towards short-lived effector cells^75^. In addition to impacts on PD1 during EBV infection, SAP regulates the SLAM-mediated organization of lytic vesicles within CD8+ T cells^76^. Thus, it is possible that SAP impacts PD1 signaling to alter CD8+ T cell differentiation into short-lived CD8 effectors and/or lytic molecule production, and/or modulating Tpex-like CD8+ T cell formation in this model. Future experiments will be required to determine which of these functions underlie the diabetes protection observed.

Overall, these data emphasize the role of SAP in fostering pro-inflammatory programming among anti-insulin B and T lymphocytes to support islet attack. Altering T-B communication, either by loss of SAP (shown here), or loss of BCL6^25^, reduces anti-insulin skewing towards atypical and GC B cell phenotypes, with some of these B cell subsets (GC B cells) noted for having features associated with enhanced antigen-presenting cell capacity^71^. Anti-insulin B cell formation did not represent a “point of no return” with respect to islet attack. Rather, our data support future studies focused on therapeutically disrupting atypical/EF and GC B:Tfh cell interactions to prevent T1D progression and diabetes onset.

## Abbreviations

(EF): Extrafollicular
(GC): germinal center
(NOD): non-obese diabetic
(ns): not significant
(pLNs): pancreatic lymph nodes
(SAP): SLAM-associated protein
(SLAM): signaling lymphocytic activation molecule
(T1D): type 1 diabetes
(Tfh): T follicular helper
(Tpex): T progenitor exhausted
(Tph): T peripheral helper
(WT): Wildtype

## Acknowledgements

This work was supported by Breakthrough T1D (formerly the Juvenile Diabetes Research Foundation) grants: 3-PDF-2024-1495-A-N (D.H.M), 2-SRA-2023-1453-S-B (R.H.B), 2-SRA-2026-1859-S-B (R.H.B.); NIH grants F30 DK145130 (L.M.C), T32 GM007347 (L.M.C, C.M.N), T32 AR059039 (D.H.M), TL1TR002244 (C.M.N.), K08 GM163052 (M.T.S), R01 AI051448 (initially awarded to Dr. James W. Thomas (Vanderbilt), transferred to R.H.B); and the American Association of Immunologists Intersect Fellowship Program for Computational Scientists and Immunologists (D.H.M).

We also acknowledge the Vanderbilt University Medical Center (VUMC) Flow Cytometry Shared Resource [supported by supported by the Vanderbilt Ingram Cancer Center (P30 CA068485) and the Vanderbilt Digestive Disease Research Center (P30 DK058404)], the VUMC Tissue Pathology Shared Resource [supported by NCI/NIH Cancer Center Support Grant, P30 CA068485], and the Islet and Pancreas Analysis Core [supported by the Vanderbilt Diabetes Research and Training Center (NIH grant P30 DK20593)].

We would like to acknowledge Dr. Pamela Schwartzberg and Dr. Jennifer Cannons (National Institutes of Health, Bethesda MD) for providing the original *SAP^-/-^*.C57BL/6 mice to Dr. James W. Thomas, which we previously backcrossed to the NOD strain ^29^ and crossed to the VH125^SD^.NOD mice (developed by Dr. James W. Thomas ^27^) for these studies. We thank Dr. James W. Thomas (Vanderbilt) for gifting the VH125^SD^.*SAP^-/-^*.NOD and control lines. We thank the NIH Tetramer Core Facility (contract number 75N93020D00005) for providing MHCI tetramers used in this study. We thank Natalie Favret and Megan Erwin (formerly Vanderbilt University) for guidance on CD8^+^ T cell staining. We thank Jason Weinstein (Rutgers New Jersey Medical School) and his lab for providing the IL-21 staining protocol, modified for this paper.

## Author contributions

Conceptualization: L.M.C, D.H.M, R.H.B. Methodology: L.M.C, D.H.M, L.E.B. Formal analysis: L.M.C., D.H.M., C.M.N, M.T.S. Investigation: L.M.C, D.H.M., J.C.M, A.M, M.L.P, C.T.B, L.E.B. Writing and editing: L.M.C, D.H.M, J.C.M, R.H.B; Funding acquisition: L.M.C., D.H.M., M.T.S., R.H.B.

## Data availability

Data acquired specifically for this study is available within the article itself and the supplemental information.

