## Supplemental Figures for "SAP loss limits anti-insulin atypical B cell activation and pro-inflammatory CD8 T cells despite preserved Tfh responses to protect against type 1 diabetes"

### SUPPLEMENTAL MATERIALS

| Supplemental Table 1: Flow Cytometry Antibody and Clones | | | | | | |
|---|---|---|---|---|---|---|
| **Antigen** | **Fluorochrome** | **Company/Clone** | **Identifier** | **Dilution** | **Conc.** | **Final Dil. ug/ml** |
| CXCR5 | BV421 | BDBioscience (2G8) | 562889 | 1:50 | 0.2 mg/ml | 4 |
| ICOS | Pacific Blue | BioLegend (C398.4) | 313521 | 1:250 | 0.5 mg/ml | 2 |
| CD23 | BV510 | BioLegend (B3B4) | 101623 | 1:125 | 0.2 mg/ml | 1.6 |
| CD69 | Superbright Violet 600 | Thermofisher (H1.2F3) | 63-0691-82 | 1:250 | 0.2 mg/ml | 0.8 |
| CD19 | Superbright Violet 645 | Thermofisher (1D3) | 64-0193-82 | 1:125 | 0.2 mg/ml | 1.6 |
| Ki67 | BV711 | Thermofisher (SolA15) | 407-5698-82 | 1:1600 | 0.2 mg/ml | 0.125 |
| IgD | BV750 | BDBiosciences (217-170) | 746873 | 1:250 | 0.2 mg/ml | 0.8 |
| CD4 | Super Bright 780 | Thermofisher (GK1.5) | 78-0041-82 | 1:500 | 0.2 mg/ml | 0.4 |
| GL7 | FITC | BDBioscience (GL7) | 553666 | 1:100 | 0.5 mg/ml | 5 |
| CD45 | SparkBlue574 | BioLegend (30-F11) | 103183 | 1:500 | 0.5 mg/ml | 1 |
| CD95 (Fas) | RB744 | BDBioscience (Jo2) | 756863 | 1:125 | 0.2 mg/ml | 1.6 |
| Igk light chain | RB780 | BDBiosciences (187.1) | 755937 | 1:250 | 0.2 mg/ml | 0.8 |
| PD-1 | PE | BioLegend (RMP1-30) | 109103 | 1:50 | 0.2 mg/ml | 4 |
| Streptavidin | PE-Dazzle594 | BioLegend | 405247 | 1:1000 | 0.2 mg/ml | 0.2 |
| Human insulin | Biotin | Sigma-Aldrich | I2643 | 1:112, then 1:100 | N/A | N/A |
| IgM | PE-Cy5 | BioLegend (RMM-1) | 406543 | 1:125 | 0.2 mg/ml | 1.6 |
| CD44 | PE-Fire810 | BioLegend (IM7) | 103085 | 1:500 | 0.2 mg/ml | 0.4 |

| | | | | | | |
|---|---|---|---|---|---|---|
| CD86 | APC-R700 | BDBioscience (GL1) | 566479 | 1:1000 | 0.2 mg/ml | 0.2 |
| Viability Dye | Viakrome808 | Beckman Coulter | C36628 | 1:1000 | N/A | N/A |
| CD21/35 | APC-Fire750 | BioLegend (7E9) | 123433 | 1:500 | 0.2 mg/ml | 0.4 |
| CD138 | APC | BioLegend (281-2) | 142505 | 1:400 | 0.2 mg/ml | 0.5 |
| TACI(CD267) | BV421 | BD Biosciences (8F10) | 742840 | 1:125 | 0.2 mg/ml | 1.6 |
| CD80 | RB705 | BDBioscience (B7-1) | 570553 | 1:250 | 0.2 mg/ml | 0.8 |
| CD11c | Pacific Blue | BioLegend (N418) | 117321 | 1:500 | 0.5 mg/ml | 1 |
| B220 | RB545 | BDBioscience (RA3-6B2) | 756542 | 1:100 | 0.2 mg/ml | 2 |
| PDL2 | RB613 | BDBioscience (MIH37) | 758326 | 1:125 | 0.2 mg/ml | 1.6 |
| CD73 | PercpCy5.5 | BioLegend (TY/11.8) | 127213 | 1:500 | 0.2 mg/ml | 0.4 |
| T-bet | PE | BioLegend (4B10) | 644809 | 1:250 | 0.2 mg/ml | 0.8 |
| CD38 | RY703 | BDBioscience (90/CD38) | 771651 | 1:4000 | 0.2 mg/ml | 0.05 |
| CD11b | APC-Fire810 | BioLegend (M1/70) | 101261 | 1:250 | 0.2 mg/ml | 0.8 |
| CXCR3 | BV480 | Invitrogen (CXCR3-173) | 414-1831-82 | 1:125 | 0.2 mg/ml | 1.6 |
| CD62L | BV570 | BioLegend (MEL-14) | 104433 | 1:500 | 25 ug/ml | 0.05 |
| CD3 | RB780 | BDBiosciences (17A2) | 755792 | 1:125 | 0.2 mg/ml | 1.6 |
| CCR6 | PE-Fire700 | BioLegend (29-2L17) | 129836 | 1:125 | 0.2 mg/ml | 1.6 |
| Foxp3 | PE-Cy7 | Invitrogen (FJK-16s) | 25-5773-80 | 1:4000 | 0.2 mg/ml | 0.05 |
| Bcl6 | Alexa 647 | BDBiosciences (K112-91) | K112-91 | 1:100 | 100 ug/ml | 1 |
| CD8a | APC-Fire750 | BioLegend (53-6.7) | 100765 | 1:500 | 0.2 mg/ml | 0.4 |
| CD25 | APC-Fire810 | BioLegend (PC61) | 102075 | 1:500 | 0.2 mg/ml | 0.4 |

| | | | | | | |
|---|---|---|---|---|---|---|
| CD11c | PacBlue | BioLegend (N418) | 117321 | 1:500 | 0.5 mg/ml | 1 |
| CD23 | BV510 | BioLegend (B3B4) | 101623 | 1:125 | 0.2 mg/ml | 1.6 |
| CD19 | SBV645 | Invitrogen (eBio1D3 (1D3)) | 64-0193-82 | 1:125 | 0.2 mg/ml | 1.6 |
| IgD | BV750 | BDBioscience (AMS 9.1) | 747077 | 1:250 | 0.2 mg/ml | 0.8 |
| HINS | FITC | Sigma-Aldrich | I2643 | 1:400 surface, 1:3200 intracellular | N/A | N/A |
| B220 | RB545 | BDBioscience (RA3-6B2) | 756542 | 1:100 | 0.2 mg/ml | 2 |
| PDL2 | RB613 | BDBioscience (MIH37) | 758326 | 1:125 | 0.2 mg/ml | 1.6 |
| CD73 | PercpCy5.5 | BioLegend (TY/11.8) | 127213 | 1:500 | 0.2 mg/ml | 0.4 |
| CD80 | RB705 | BD Bioscience (16-10A1) | 570553 | 1:250 | 0.2 mg/ml | 0.8 |
| CD95 | RB744 | BDBioscience (JO2) | 756863 | 1:125 | 0.2 mg/ml | 1.6 |
| Igk | RB780 | BDBioscience (187.1) | 755937 | 1:250 | 0.2 mg/ml | 0.8 |
| Blimp1 | PE-CF594 | BDBioscience (5E7) | 564269 | 1:1600 | 0.2 mg/ml | 0.125 |
| T-bet | PE | BioLegend (4B10) | 644809 | 1:250 | 0.2 mg/ml | 0.8 |
| IgM | PE-Cy5 | BioLegend (RMM-1) | 406543 | 1:125 | 0.2 mg/ml | 1.6 |
| CD38 | RY703 | BDBioscience (90/CD38) | 771651 | 1:4000 | 0.2 mg/ml | 0.05 |
| IgG2a-bio | PE-Cy7 | BDBioscience (8.3) | 553502 | 1:2000 biotin, 1:100 Streptavidin | 0.5 mg/mL | 0.25 |
| CD44 | PE-Fire810 | BioLegend (IM7) | 103085 | 1:500 | 0.2 mg/ml | 0.4 |
| CD138 | APC | BioLegend (281-2) | 142505 | 1:400 | 0.2 mg/ml | 0.5 |
| CD86 | APC-R700 | BDBioscience (GL1) | 565479 | 1:1000 | 0.2 mg/ml | 0.2 |

| | | | | | | |
|---|---|---|---|---|---|---|
| CD21 | APC-Fire750 | BioLegend (7E9) | 123433 | 1:500 | 0.2 mg/ml | 0.4 |
| CD11b | APC-Fire810 | BioLegend (M1/70) | 101288 | 1:250 | 0.2 mg/ml | 0.8 |
| H-2K$^d$/NRP-V7 (KYNKANVFL) | PE | NIH Tetramer Core | N/A | 1:100 | 1.33 mg/ml | 13.3 |
| H-2K$^d$/Insulin B chain 15-23 (LYLVCGERL) | BV421 | NIH Tetramer Core | N/A | 1:100 | 1.17 mg/ml | 11.7 |

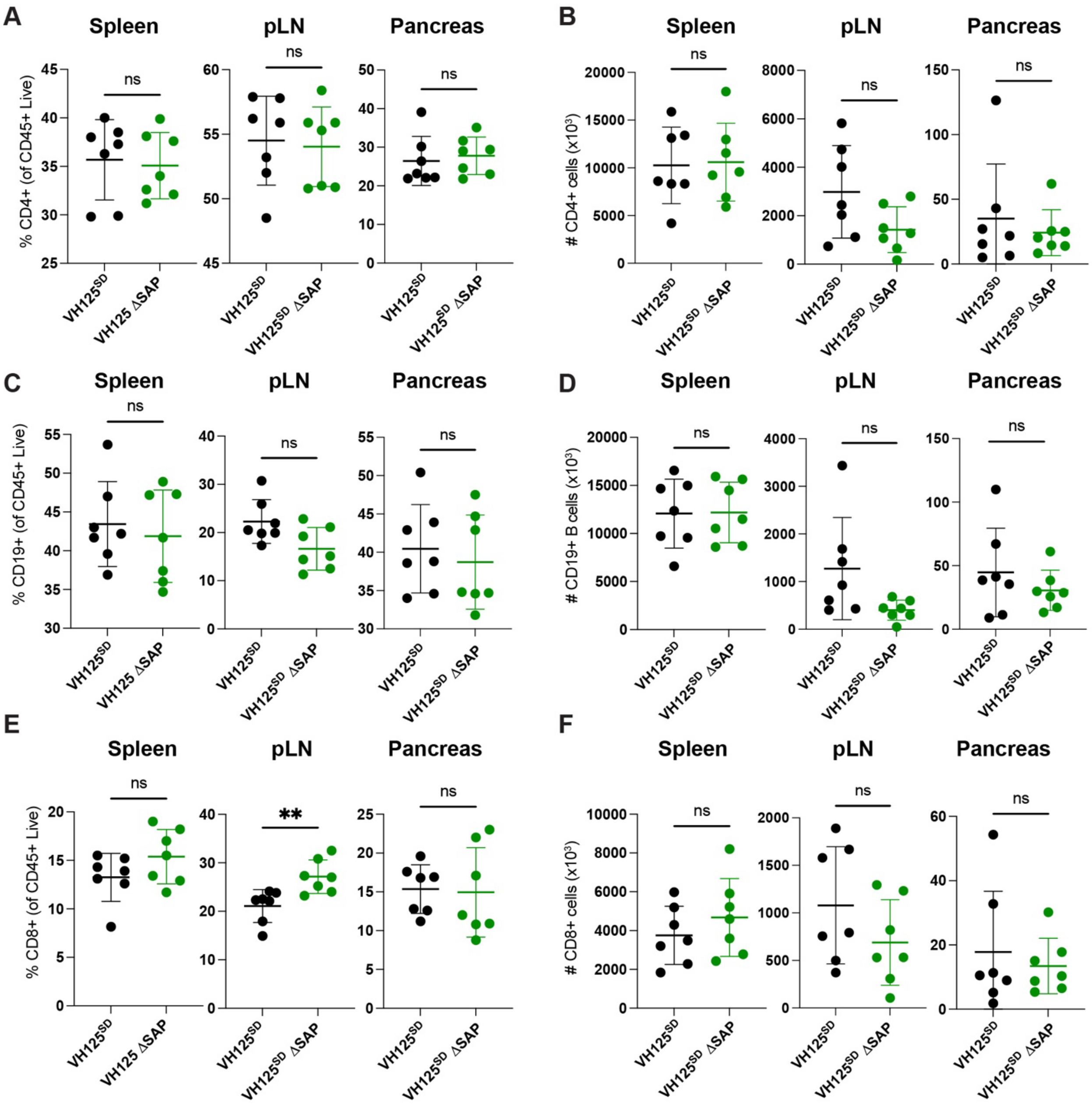


**Supplemental Figure 1: CD4, CD8, and CD19+ populations are not altered by loss of SAP in VH125$^{SD}$ mice.** Cells from the indicated organs were isolated from 12-14-wk-old, prediabetic VH125$^{SD}$ mice with and without SAP and were subjected to flow cytometry analysis. **(A)** Proportion and **(B)** number of CD4+ T cells, **(C)** proportion and **(D)** number of CD19+ cells, and **(E)** proportion and **(F)** number of CD8+ cells in pLNs. For all experiments, n = 6-8 mice per group, bars represent mean +/- SD. ** $p < 0.01$, Mann-Whitney U-test.

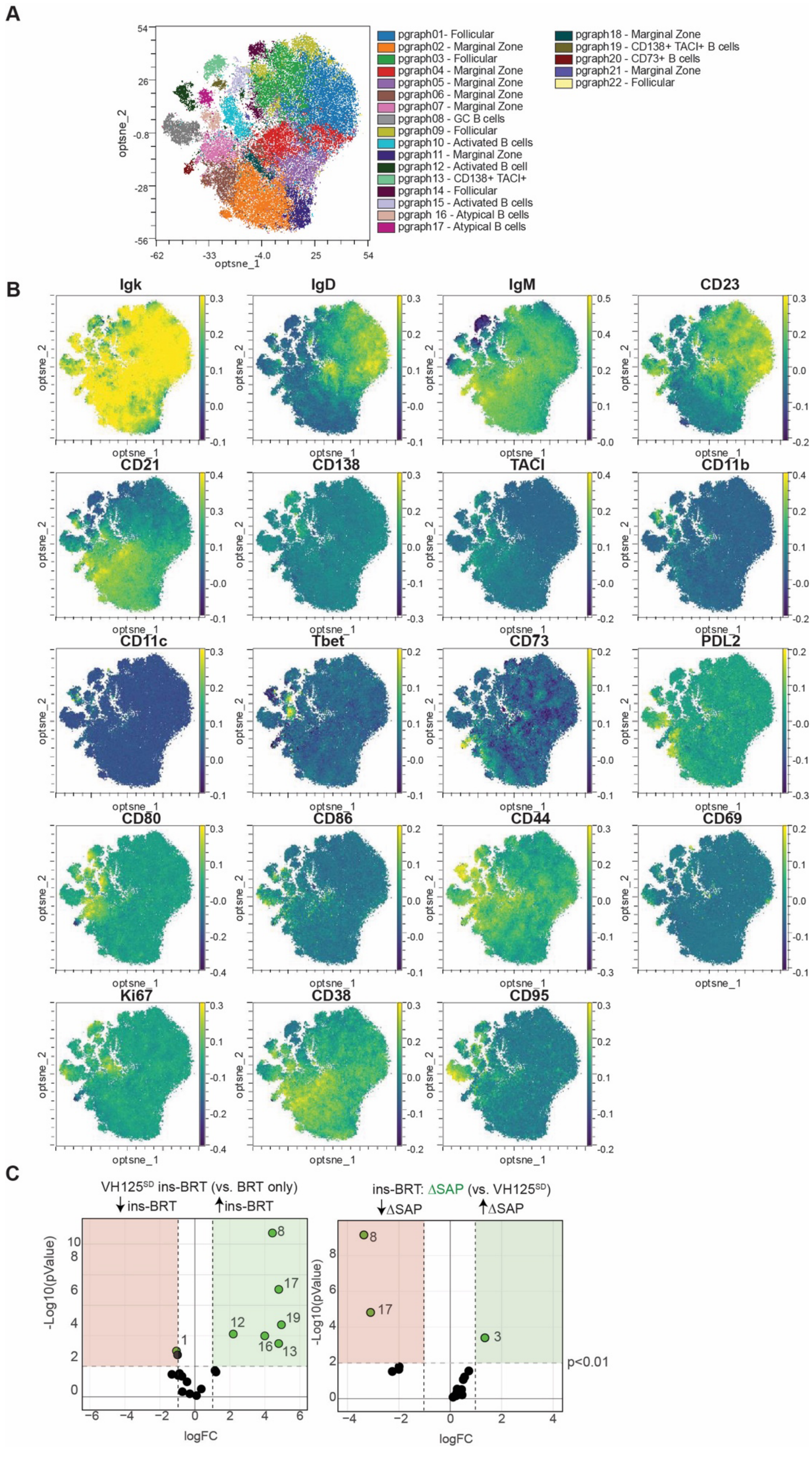

A
optsne_2
optsne_1
54
26
-0.8
-28
-56
-62
-33
-4.0
25
54
pgraph01- Follicular
pgraph02 - Marginal Zone
pgraph03 - Follicular
pgraph04 - Marginal Zone
pgraph05 - Marginal Zone
pgraph06 - Marginal Zone
pgraph07 - Marginal Zone
pgraph08 - GC B cells
pgraph09 - Follicular
pgraph10 - Activated B cells
pgraph11 - Marginal Zone
pgraph12 - Activated B cell
pgraph13 - CD138+ TACI+
pgraph14 - Follicular
pgraph15 - Activated B cells
pgraph 16 - Atypical B cells
pgraph17 - Atypical B cells
pgraph18 - Marginal Zone
pgraph19 - CD138+ TACI+ B cells
pgraph20 - CD73+ B cells
pgraph21 - Marginal Zone
pgraph22 - Follicular
B
Igk
IgD
IgM
CD23
CD21
CD138
TACI
CD11b
CD11c
Tbet
CD73
PDL2
CD80
CD86
CD44
CD69
Ki67
CD38
CD95
optsne_1
optsne_2
C
VH125SD ins-BRT (vs. BRT only)
↓ins-BRT
↑ins-BRT
-Log10(pValue)
logFC
ins-BRT: ΔSAP (vs. VH125SD)
↓ΔSAP
↑ΔSAP
p<0.01

**Supplemental Figure 2: Distinct subsets of anti-insulin B cells that expand following insulin-BRT immunization and depend on SAP.** Anti-insulin B cells (CD45+ CD19+ B220+ INS+ live singlets) were subsampled following insulin-BRT or BRT-only immunization in VH125$^{SD}$ and VH125$^{SD}$ΔSAP mice. **(A)** Phenograph clustering algorithm identified n = 22 distinct clusters of anti-insulin B cells. The indicated population names were assigned based on **(B)** individual marker expression plots of spectral flow cytometry panel. **(C)** Differential abundance analysis via diffcyt for clustered populations was used to compare either ins-BRT vs. BRT-only immunization in VH125$^{SD}$ SAP-sufficient mice (*left*), or ins-BRT immunization comparing SAP-sufficient and deficient VH125$^{SD}$ mice (*right*). Green dots represent p value < 0.01 and |log2FC| > 1, with clusters identities labeled.

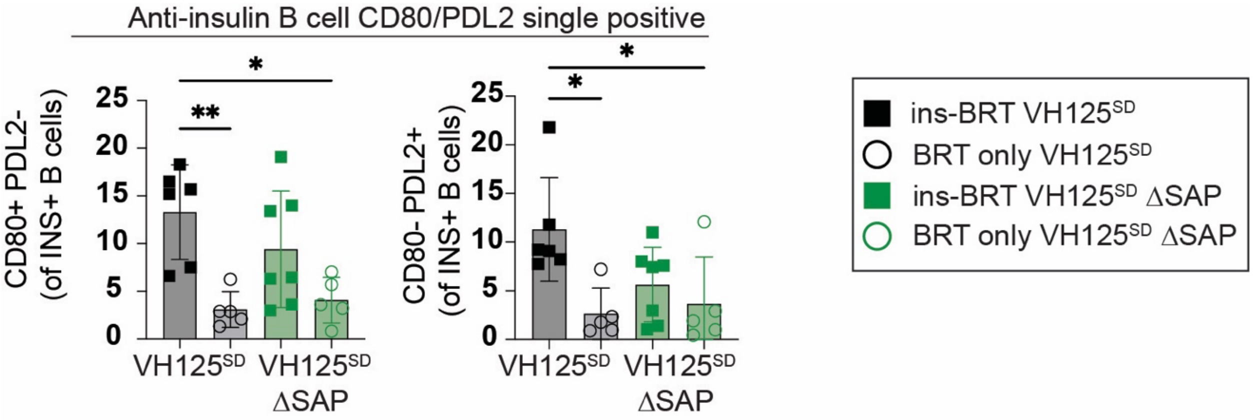


**Supplemental Figure 3: A memory-like anti-insulin B cell subset expands following insulin-BRT immunization.** The proportions of CD80+ PDL2- (*Left*) and CD80- PDL2+ (*Right*) are shown amongst insulin-binding B cells. n = 5-7 mice per group, error bars represent mean +/- SD. * $p < 0.05$, ** $p < 0.01$, one-way ANOVA and post-hoc Tukey’s multiple comparisons test.

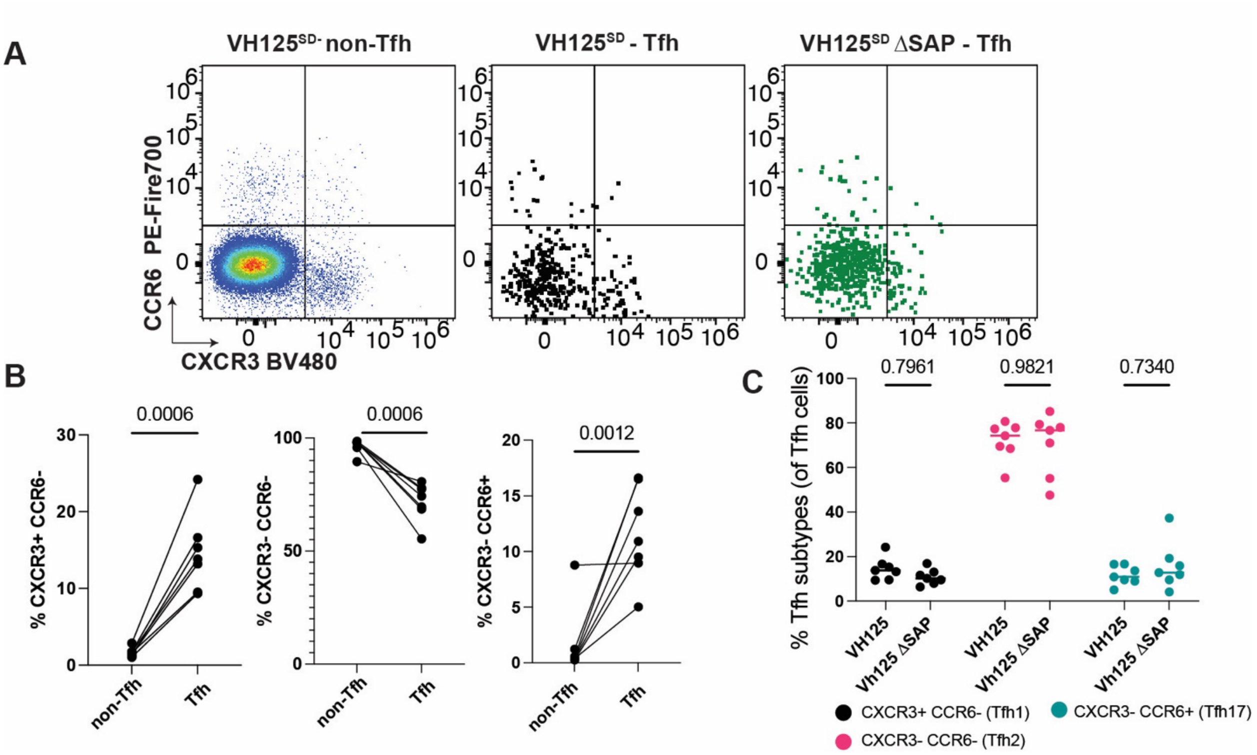


**Supplemental Figure 4: Loss of SAP does not impact Tfh differentiation, proliferation, or activation.** pLNs tissues were harvested from 12-16-week-old, prediabetic VH125$^{SD}$ and VH125$^{SD}\Delta$SAP mice, n = 6-8 mice per group are plotted. **(A)** Representative flow plots show expression of CCR6 and CXCR3, which identifies CXCR3+ (Th1/Tfh1-like), CXCR3-CCR6- (Th2/Tfh2-like), and CCR6+ populations (Th17/Tfh17-like), amongst non-Tfh, Tfh, and Tfh-ΔSAP groups, with quantification of **(B)** non-Tfh and Tfh comparisons and **(C)** % Tfh1, Tfh2, or Tfh17-like populations. All were ns by *(B)* Mann Whitney U test or *(C)* Kruskal-Wallis test with post-hoc Dunn’s test.

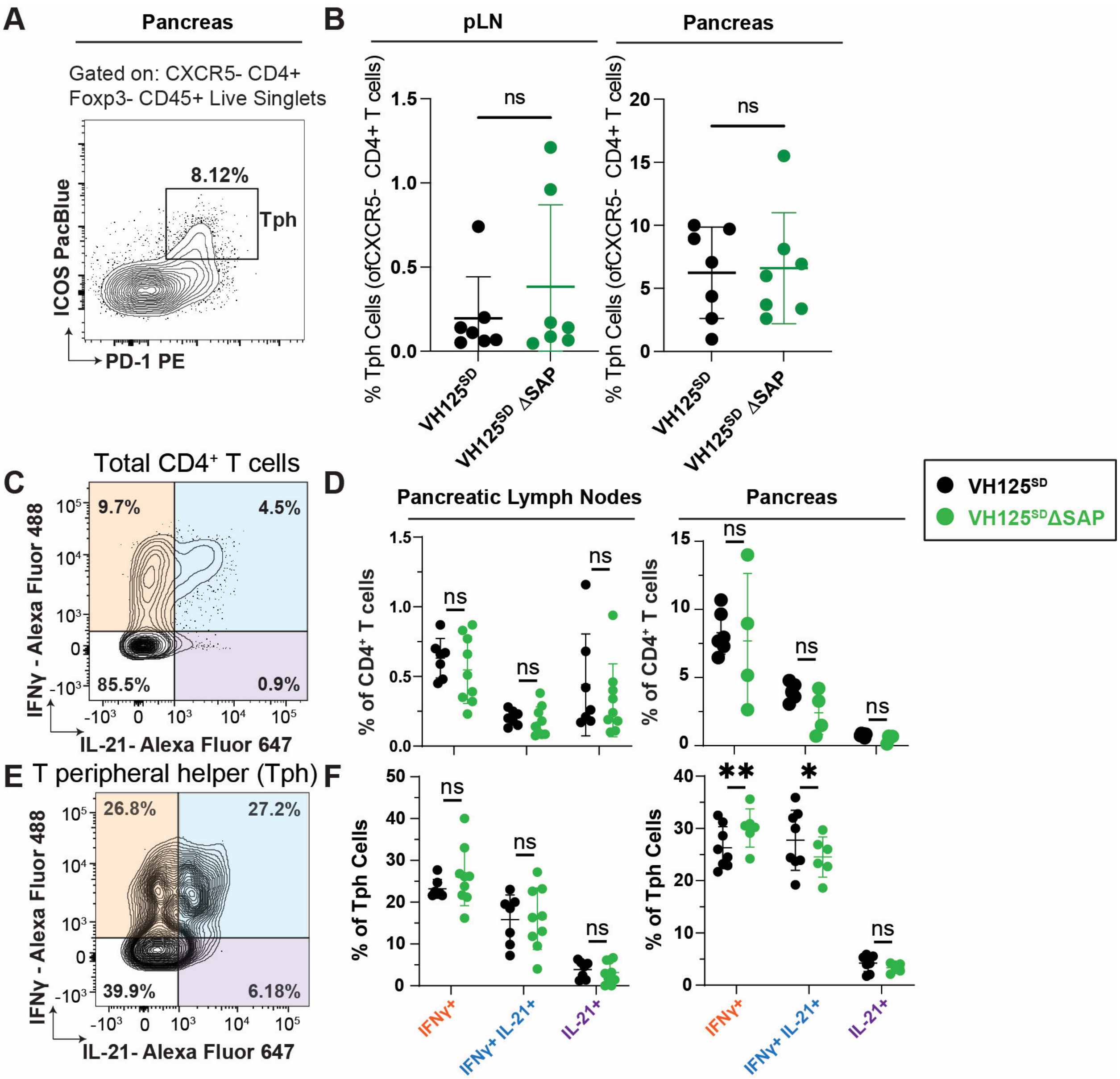


**Supplemental Figure 5: T peripheral helper cells are not altered in proportion or in IFN-γ/IL-21 cytokine skewing with the loss of SAP.** Pancreas and pLNs were harvested from 12-16-wk-old, prediabetic VH125$^{SD}$ (black) and VH125$^{SD}$ΔSAP mice (green), n = 6-9 mice per group. **(A, C)** Representative plot and scheme to identify **(A)** T peripheral helper (Tph) cells (CXCR5- ICOS+ PD-1+ CD4+ Foxp3- CD45+ live lymphocytes) and **(C, E)** IL-21 and IFN-γ staining in **(C)** total CD4+ or **(E)** Tph cells. **(B)** % Tph (of CXCR5- CD4+ T cells) in pLNs (*left*) and pancreas (*right*). **(D, F)** % IFN-γ+, IFN-γ+ IL-21+, and IL-21+ subsets of **(D)** total CD4+ or **(F)** Tph cells. Bars represent mean +/- SD. * $p < 0.05$, ** $p < 0.01$, Mann Whitney U test.

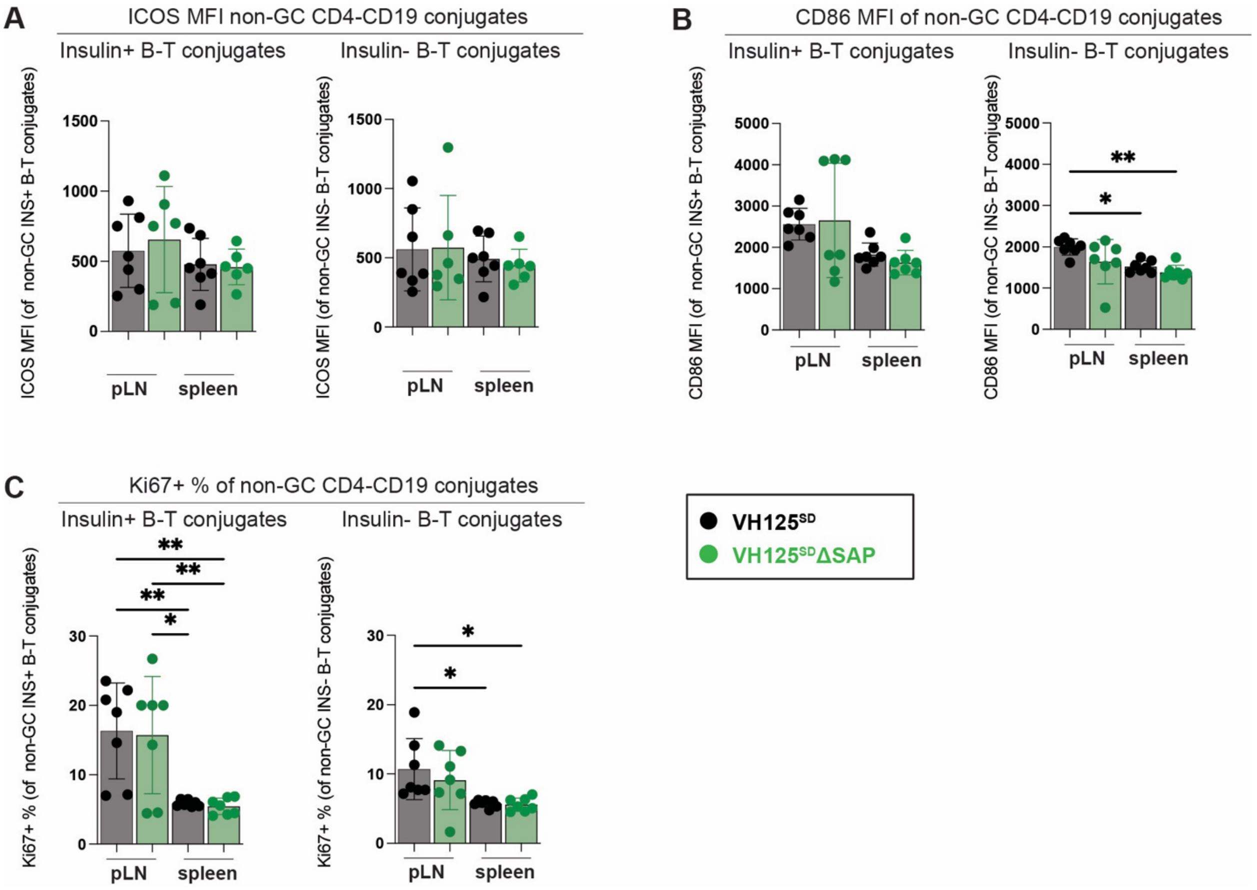


**Supplemental Figure 6: Anti-insulin B-CD4+ T cell conjugates formed *in vivo* show increased ICOS, CD86, and % Ki67+ levels in pLNs vs. spleen, which are not altered by loss of SAP.** CD4+ T-B conjugates were identified among either insulin+ or insulin- populations as in **Figure 6A** in the pLNs or spleen of 12-16-wk-old, prediabetic VH125$^{SD}$ and VH125$^{SD}$ ΔSAP mice. **(A)** ICOS MFI, **(B)** CD86 MFI, and **(C)** % Ki67+ is shown for n = 6-8 mice per group. Bars represent mean +/- standard deviation, * p ≤ 0.05, ** p ≤ 0.01, one-way ANOVA with post-hoc Tukey's multiple comparison test.
